# THE WIP6 TRANSCRIPTION FACTOR *TOO MANY LATERALS* SPECIFIES VEIN TYPE IN C_4_ AND C_3_ GRASS LEAVES

**DOI:** 10.1101/2023.12.20.572592

**Authors:** Daniela Vlad, Maricris Zaidem, Chiara Perico, Olga Sedelnikova, Samik Bhattacharya, Jane A. Langdale

## Abstract

Grass leaves are invariantly strap shaped with an elongated distal blade and a proximal sheath that wraps around the stem. Underpinning this uniform shape is a scaffold of leaf veins, most of which extend in parallel lines along the proximo-distal leaf axis. Differences between species are apparent both in the types of veins that develop and in the spacing between them across the medio-lateral leaf axis. A prominent engineering goal is to increase vein density and the proportion of bundle sheath cells surrounding the veins in leaves of C_3_ photosynthesizing species such as rice, in order to facilitate introduction of the more efficient C_4_ photosynthetic pathway. Here we discover that the WIP6 zinc finger transcription factor TOO MANY LATERALS (TML) specifies vein rank in both maize (C_4_) and rice (C_3_), species with distinct venation patterns. Loss of function *tml* mutations lead to the development of large lateral veins in positions normally occupied by smaller intermediate veins. The spatial localization of *TML* transcripts in wild-type leaves is consistent with a role in suppressing lateral vein formation in procambial cells that develop intermediate veins, specifically the class of intermediate veins that extend from the leaf blade into the leaf sheath. Attempts to manipulate TML function in rice were unsuccessful because transgene expression was silenced, suggesting that precise spatial and temporal regulation of *TML* expression is essential during the regeneration of shoot tissue from callus. Given that transcriptome analysis demonstrated altered profiles of genes associated with cytokinin and auxin signaling in loss of function maize mutants, the necessity for tight regulation of *TML* gene expression could be an indirect consequence of hormonal inbalances as opposed to ectopic activity of a specific downstream target. Importantly, however, loss of function mutants in rice display increased vascular and bundle sheath cell occupancy in the leaf. Collectively this work provides an understanding of how vein rank is specified in grass leaves and a first step towards an anatomical chassis for C_4_ engineering in rice.

## INTRODUCTION

Biological patterns form during the development of multicellular organisms when cell fates are mapped onto fields of equivalent pluripotent cells in organized space and time ^1^. In flowering plants, patterning primarily occurs post-embryogenesis at the shoot and root apical meristems, with shoot patterns most obvious in the geometrical arrangement of leaves around the apex and in floral morphologies ^2^. Less obvious is the patterning of veins within leaves, which in flowering plants is of two distinct types. In eudicotyledonous (eudicot) plants, leaf veins form reticulated networks whereas in monocotyledonous (monocot) plants, veins run in parallel along the length of the proximo-distal leaf axis (reviewed in ^3,4,5^). Both the reticulate venation pattern in eudicot leaves and the striated pattern in monocot leaves result from the organized specification of vein-forming stem cells (procambial initial cells) within the ground meristem of the leaf ^6^, with temporal regulation facilitating the sequential formation of different vein ranks and spatial regulation defining the distance between veins.

Vein patterning mechanisms are best characterized in eudicots where the regulated flux of auxin determines the direction and extent of vein development. In the eudicot *Arabidopsis thaliana* (Arabidopsis), auxin flux occurs both via passive diffusion through plasmodesmata ^7^ and active transport through membrane localized transporters ^8,9^, with the asymmetric distribution of efflux carriers such as PIN FORMED1 (PIN1) directing flux from the leaf margin to the midvein through narrow strands of procambium (reviewed in ^10^). The specification of procambial initials (referred to as preprocambial cells in Arabidopsis ^11^) that form the procambial strands occurs when auxin signaling activates the expression of the auxin response factor *MONOPTEROS/ARF5* (*MP*) ^12^. MP then elevates PIN1 expression in a feedback loop and induces expression of the HD-ZIP III transcription factor *ARABIDOPSIS THALIANA HOMEOBOX 8* (*ATHB8*), which marks procambium identity and stabilizes PIN1 localization ^13^. The PIN/MP/ATHB8 module thus reinforces auxin flux and leads to the extension of procambial strands. Although procambium develops simultaneously along a file of preprocambial cells, it is important to note that veins form sequentially in rank order with new procambial strands developing towards, and connecting with, those that have already formed ^14,15^.

Little is known about vein patterning in monocot leaves but mechanisms appear inherently different from those in eudicots because successive vein ranks are positioned in between and parallel to existing veins, with connections only made late in development when transverse veins form across the medio-lateral leaf axis. Vein ontogeny in monocots is best characterized in grasses such as *Zea mays* (maize) where four longitudinal vein ranks develop ^16,17^. The midvein develops first, differentiating from the base of the leaf primordium to the tip, the lateral veins develop next (also towards the leaf tip) and then the rank 1 and rank 2 intermediate veins develop in turn from the tip towards the base. Only the largest of the rank 1 veins, which are associated with sclerenchyma on both the adaxial and abaxial sides, extend into the leaf sheath and as such the veins in the sheath are spaced further apart than those in the blade. In maize, the four longitudinal vein ranks underpin a cellular arrangement known as Kranz anatomy in which each vein is surrounded by concentric rings of photosynthetic bundle sheath cells and mesophyll cells, and all veins in the blade are separated by just four cells in the medio-lateral leaf axis (reviewed in ^18^). This anatomy, which is associated with C_4_ photosynthesis, evolved on multiple independent occasions from the ancestral anatomy found in C_3_ photosynthesising grasses such as *Oryza sativa* (rice) ^19^. In C_3_ grass leaves, rank 2 intermediate veins are absent and vein pairs in the blade are separated by up to ten mesophyll cells. Venation patterns in C_4_ grass leaves therefore differ from those in C_3_ grasses, both in terms of the spacing between veins (closer in C_4_ than C_3_) and in the number of vein ranks (more in C_4_ than C_3_) (reviewed in ^3^). In both C_4_ and C_3_ leaves, however, the timing of vein initiation is similar. The midvein initiates at plastochron (P) 1 (where a plastochron is the time interval between initiation of primordia at the meristem and P1 is the most recently formed), the laterals at P2/P3 and the intermediates between P3 and P5 ^16,20,21^. In each case, as in eudicots, cells in procambial strands are marked by expression of a *PIN* gene ^22–25^ but the strands are not directed towards existing veins, suggesting that the spacing of veins across the medio-lateral leaf axis is pre-patterned within ground meristem cells of the emerging primordium. With the exception of the homeobox gene *OsHOX1,* which is expressed in procambial cells of developing rice leaf primordia ^26^, the identity of genes acting downstream of auxin during the formation and extension of procambial strands in monocot leaves remains elusive.

Here we identify a gene encoding a WIP C2H2 zinc finger protein ^27^ that is expressed in procambial initial cells in the leaves of multiple grass species. The maize gene Zm00001d020037 (B73 v3 - GRMZM2G150011; B73 v5 - Zm00001eb309530) was first identified as a candidate regulator of vein patterning in a transcriptome analysis of maize foliar and husk leaf primordia ^28^. Transcript accumulation profiles from P1 to P5 indicated an increase in expression levels prior to the initiation of veins in developing P1/P2 primordia and a decrease after P5 once all veins had been specified. The gene was hypothesized to be a regulator of Kranz anatomy ^29^. Through loss of function analyses in maize and *Setaria viridis* we demonstrate that the WIP6 gene *TOO MANY LATERALS* (*TML)* facilitates development of the correct ratio of lateral to intermediate veins in C_4_ grass leaves. Rather than being specific for Kranz anatomy, however, we show that aspects of TML function are conserved between C_4_ and C_3_ species, enabling venation patterns to be modified in rice.

## RESULTS

### A canonical EAR domain is present in monocot but not eudicot WIP6 proteins

As a first step towards understanding the likely function of the maize gene Zm00001d020037, orthologous genes were identified in predicted proteomes of all species using Orthofinder ^30^ (**Supplementary File 1**). The orthogroup contains WIP proteins, which are specific to land plants and are characterized by a conserved three amino acid ‘WIP’ sequence and four zinc finger domains ^31^. Phylogenetic inference revealed a single gene in the bryophyte *Marchantia polymorpha* (*MpWIP*), evidence of a gene duplication that generated WIP6 and a second clade in the last common ancestor of vascular plants, and further duplications in the last common ancestor of the flowering plants which subdivided the second clade into WIP1, WIP3 and WIP2/4/5 clades (**Figure 1A, Supplementary Files 2 & 3**). Zm00001d020037 and its paralog Zm00001d005566 are members of the WIP6 clade. The only gene in this clade for which function has been reported is Arabidopsis *WIP6,* which was named *DEFECTIVELY ORGANISED TRIBUTARIES 5* (*DOT5*) on the basis of leaf venation defects observed in a T-DNA insertion line ^32^. However, recent work refuted a role for *AtWIP6* in venation patterning, being unable to identify any developmental perturbations in loss of function mutants created by gene editing ^33,34^. As such, in the absence of a known function, we initially named the maize genes *ZmWIP6A* (Zm00001d020037) and *ZmWIP6B* (Zm00001d005566).

**Figure 1.**
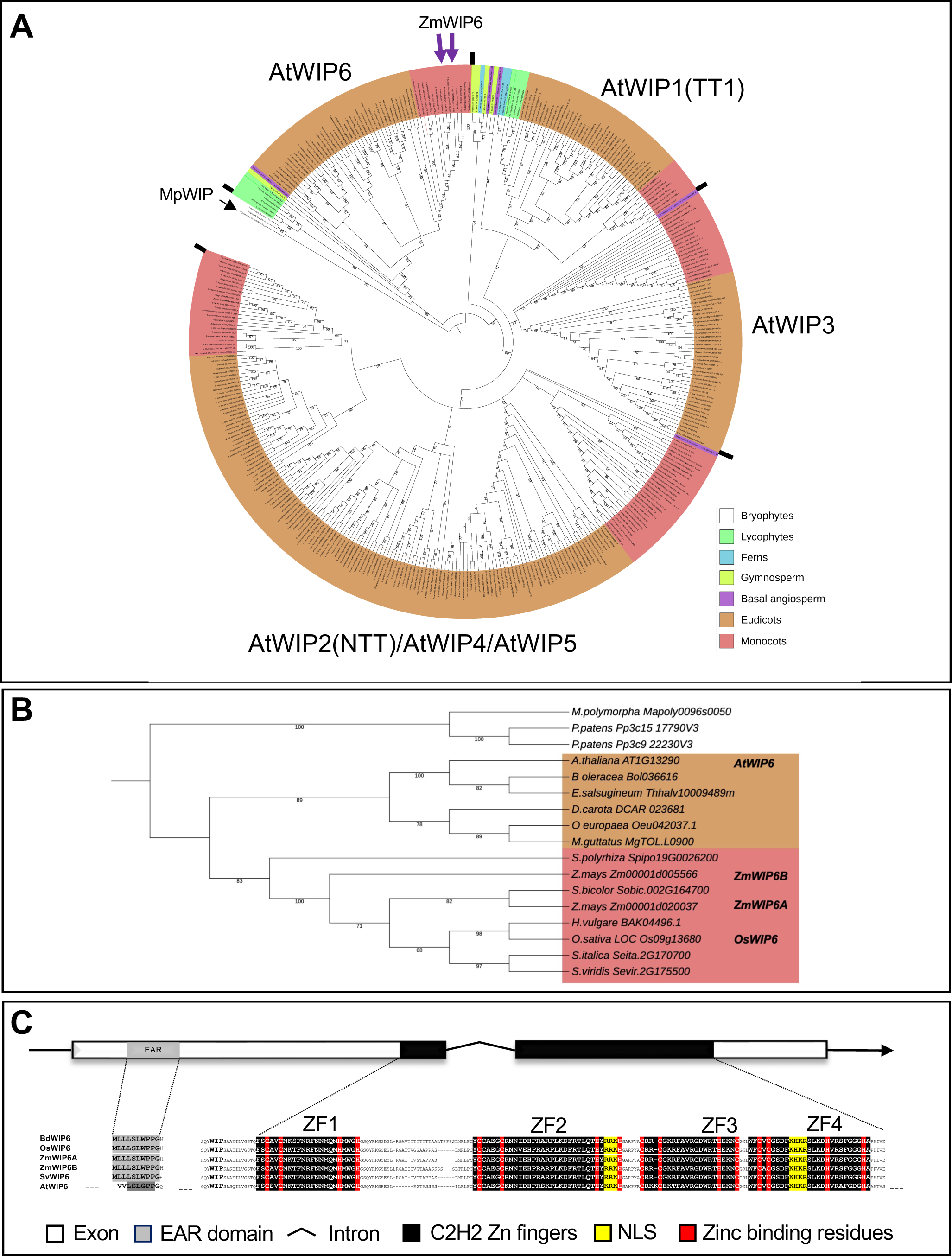
Monocot WIP6 orthologs contain a conserved N terminal EAR domain. **A)** Maximum Likelihood phylogeny of WIP proteins in land plants, rooted with the single WIP gene from *Marchantia polymorpha* (*MpWIP*). Six WIP genes are present in *Arabidopsis thaliana*, with four clades identified in basal angiosperms (WIP1, WIP3, WIP2,4,5 and WIP6), and two clades (WIP6 and WIP1-5) identified in all vascular plant lineages. Maize has two genes in the WIP6 clade. **B)** Maximum likelihood phylogeny of WIP6 proteins from representative eudicots (orange shading) and monocots (pink shading), rooted with the bryophyte clade. Bootstrap values are indicated below branches. **C)** Schematic representation of monocot WIP6 genes showing 5’ and 3’ UTRs (black lines) plus two exons (rectangles) separated by an intron (^). The arrow indicates the direction of transcription, the conserved EAR domain (grey box) and the C2H2 zinc finger region (black boxes) are highlighted. The protein alignment illustrates the amino acid residues in the conserved protein domains. The LxLxL type EAR motif present in monocot WIP6 proteins is not present in eudicot proteins (Figure S1). A WIP domain (in bold) and up to four zinc fingers (ZF) are predicted. The ZFs are highlighted in black with the zinc binding residues in red and the nuclear localization signal (NLS) is highlighted in yellow.

Within the WIP6 clade, eudicot and monocot sequences form two discrete sub-clades (**Figure 1A, 1B, & S1A, Supplementary Files 4-6**). In the monocot clade, with the exception of maize, most genomes contain a single copy of WIP6 as seen in the basal monocots *Zostera marina* and *Spirodela polyrhiza*. In maize, *ZmWIP6A* (chromosome 7) and *ZmWIP6B* (chromosome 2) are not in syntenic regions of the genome ^35^ and are thus unlikely to have arisen from the whole genome duplication event that generated many homeologous genes in the species. A protein alignment of 50 predicted WIP6 protein sequences from 41 species highlights the WIP sequence plus the four highly conserved zinc finger domains arranged in tandem at the C terminus ^27,31,36^ (**Figure 1C, Figure S1B, Supplementary File 7**). Towards the N terminus, a putative EAR motif (LxLxPP) ^37^ is present in both eudicot and bryophyte proteins but only the monocot proteins contain a *bona fide* LxLxL-type EAR motif ^38^. Given that LxLxL EAR motifs are known to mediate transcriptional repression of target genes ^37^, the observed monocot/eudicot differences in this region could reflect a repressor role for WIP6 proteins specifically in monocots.

### *WIP6* gene transcripts accumulate in procambial cells that will form intermediate veins in grass leaf primordia

In the context of whole leaf primordia, transcripts of both *ZmWIP6A* and *ZmWIP6B* accumulate to relatively low levels (550 and 80 transcripts per million respectively ^28^), but expression analyses at the whole tissue level can mask big differences between individual cells within the tissue. To evaluate the spatial pattern of *ZmWIP6A* and *ZmWIP6B* expression, *in situ* hybridization experiments were performed with sections of maize shoot apices that included late P2-5 leaf primordia. For orientation, Figure 2A depicts a schematic of a P4 leaf primordium showing the position of procambial initial cells and of procambial centres (dividing procambial cells that will give rise to veins), and Figure 2B shows a schematic of lateral and intermediate veins in an expanded leaf. Transcripts were detected in developing procambial centres of P3, P4 and P5 primordia, with *ZmWIP6A* accumulating to higher levels than *ZmWIP6B* in the leaf blade (as seen in transcriptomic experiments) (**Figure 2C, 2D**). The presence of veins at different stages of development in each leaf primordium further revealed the temporal pattern of *ZmWIP6* expression, with transcripts present in the single procambial initial cell and in the first four or five derivatives of that cell during the initation of intermediate veins but absent at later stages of development (**Figure 2E**). Similar spatial expression profiles were seen in other grass species examined, with expression of the single WIP6 ortholog in each case being confined to developing veins in *Setaria viridis*, *Sorghum bicolor* and rice primordia (**Figure S2A-C**). In rice, however, where veins are initiated further apart than in the three other species, the temporal profile of *OsWIP6* expression differs in that transcripts are detectable at later stages of vein development than seen in maize (**Figure S2D**). Collectively, these data suggest that WIP6 orthologs may play a role in the specification of procambial initial cells within the ground meristem of developing leaf primordia in both C_4_ and C_3_ species.

**Figure 2.**
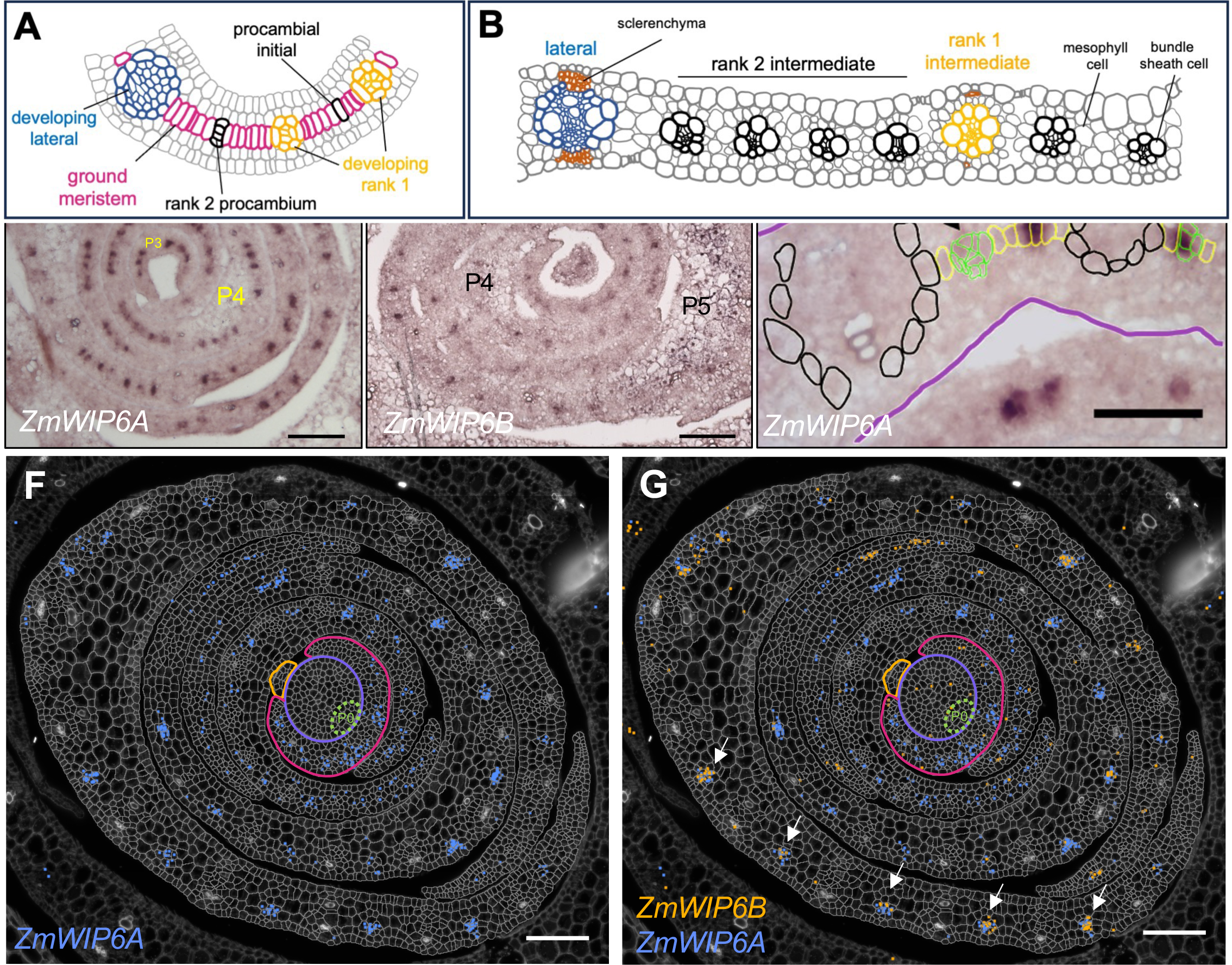
WIP6 genes are expressed in dividing procambial cells in maize leaf primordia. **A, B)** Schematics of cellular arrangements in a P4 primordium (A) and an expanded leaf (B). Different vein and cell-types are indicated. **C, D)** *In situ* hybridization of *ZmWIP6A* (Zm00001d020037) (C) and *ZmWIP6B* (Zm00001d005566) (D) transcripts to P3-P5 leaf primordia of maize. Plastochron stages are as indicated. Scale bars = 100μm **E)** Annotated image of a section hybridized with *ZmWIP6A*. The P5 primordium is outlined in purple and ground meristem cells are outlined in yellow. Four different stages of vein development are highlighted. The white arrowhead marks a procambial initial, which can only be distinguished from the surrounding ground meristem cells by the presence of *ZmWIP6A* transcripts. The green cells denote early dividing procambial centres, with the black arrowhead marking the more advanced stage. Black cells surround fully differentiated veins. *ZmWIP6A* transcripts are detected in the procambial initial and in the early dividing procambium but are absent once the developing vein comprises more than four or five cells (black arrowhead). Scale bar = 50μm. F, G) Molecular cartography showing accumulation of *ZmWIP6A* (Zm00001d020037) (blue dots in **F, G)** and *ZmWIP6B* (Zm00001d005566) (orange dots in G) transcripts in the shoot apical meristem and P1-P5 leaf primordia of maize. The purple line indicates the meristem, green dotted line the P0 primordium, orange line the P1 primordium and pink line the P2 primordium. White arrows indicate developing intermediate veins in the P5 leaf sheath tissue. Scale bar = 100μm.

To gain a more precise understanding of the spatial and temporal expression profile of *ZmWIP6* genes, and to also examine transcript accumulation patterns in the shoot apical meristem plus P0 and P1 leaf primordia, fragments of both genes were hybridized to sections of maize shoot apices using Resolve Biosciences Molecular Cartography technology. The ability to visualize individual cells and to simultaneously quantify transcripts of both genes within them revealed an overlapping but distinct expression pattern for each gene. Specifically, *ZmWIP6A* is expressed in developing intermediate veins of all leaf primordia examined, whereas *ZmWIP6B* is only expressed in a subset (**Figure 2D, E**). Neither gene is expressed in the midvein or lateral veins at any stage of development. (Note that although some signal was detected in the early P2 primordium where intermediate veins have yet to be initated, the signal was not localized to procambial centres). Co-localization of *ZmWIP6A* and *ZmWIP6B* transcripts is most apparent in developing intermediate veins of the P5 primordium (arrows in **Figure 2E & Figure S2**). In this regard it is important to note that a transverse section across the shoot apex captures leaf primordia at different positions along their proximo-distal axes. In this case, the P1 section is at the very tip of the blade, P2 and P3 are near the middle of the blade, P4 is at the base of the blade and P5 is in the leaf sheath. Given that intermediate vein development in the leaf sheath is the last stage of vein patterning within any grass leaf primordium, these observations suggest that *ZmWIP6A* may be required from the earliest stages of intermediate vein formation whereas *ZmWIP6B* function may only be required either late in development or specifically in the leaf sheath. Intriguingly, although *ZmWIP6A* activity appears to be restricted to the first few divisions in procambial centres of P3-P5 primordia, transcripts are also detected at later stages of development, alongside *ZmWIP6B* around mature veins in the leaf sheath. These observations suggest that *ZmWIP6A* may have a non-redundant specification role early in development of the leaf blade and a redundant role with *ZmWIP6B* later in the development of the leaf sheath.

### Loss of function *Zmwip6A* mutants develop too many lateral veins

To determine whether *ZmWIP6A* or *ZmWIP6B* expression patterns reflect a role in leaf venation patterning, loss of function alleles were generated by CRISPR/Cas9 mediated gene editing. Two guide RNA sequences (sgRNAs) were designed to target each gene at sites flanking the EAR domain (**Figure 3A**) and all four sgRNAs were assembled into a single construct for transformation into the maize inbred line B104. Resultant T0 plants were backcrossed to B104 and four T1 lines representing different combinations of mutant alleles were obtained (**Figure 3B**). One line contained mutations in *ZmWIP6A,* one in *ZmWIP6B,* and two in both genes. Within the four lines, there were four independent alleles of *ZmWIP6A* and three of *ZmWIP6B*, with one of the double mutant lines (438_I20) being heteroallelic at both loci. For *ZmWIP6A*, one of the mutant alleles (I20_3) encoded an almost full-length protein with just five amino acids deleted upstream of the EAR domain (**Figure S3A**). No further characterization was carried out with this allele (*Zmwip6A_m4*) because the encoded protein was likely to be at least partially functional. The remaining three *ZmWIP6A* alleles contained a 268 bp deletion (J3) or single base pair insertions (B1_10 & I20_18) that resulted in premature stop codons upstream of the zinc finger domain and thus represented loss of function alleles (**Figure S3A**). These alleles were named *Zmwip6A_m1* (J3), *Zmwip6A_m2* (B1-10) and *Zmwip6A_m3* (I20_18). For *ZmWIP6B,* a 281 bp deletion that resulted in a premature stop codon in line J3 was named *ZmWIP6B_m1.* Two further alleles were isolated from line 438_I20. One contained a 286 bp deletion (*Zmwip6B_m2*) and the second contained a 3 bp deletion plus a single bp insertion (*Zmwip6B_m3*) (**Figure S3B**). Both result in the formation of premature stop codons and predicted loss of gene function. Homozygous mutations that had segregated independently from the transgene were identified in progeny of the six self-pollinated T1 lines and single and double combinations of mutant alleles were fixed for phenotypic characterization.

**Figure 3.**
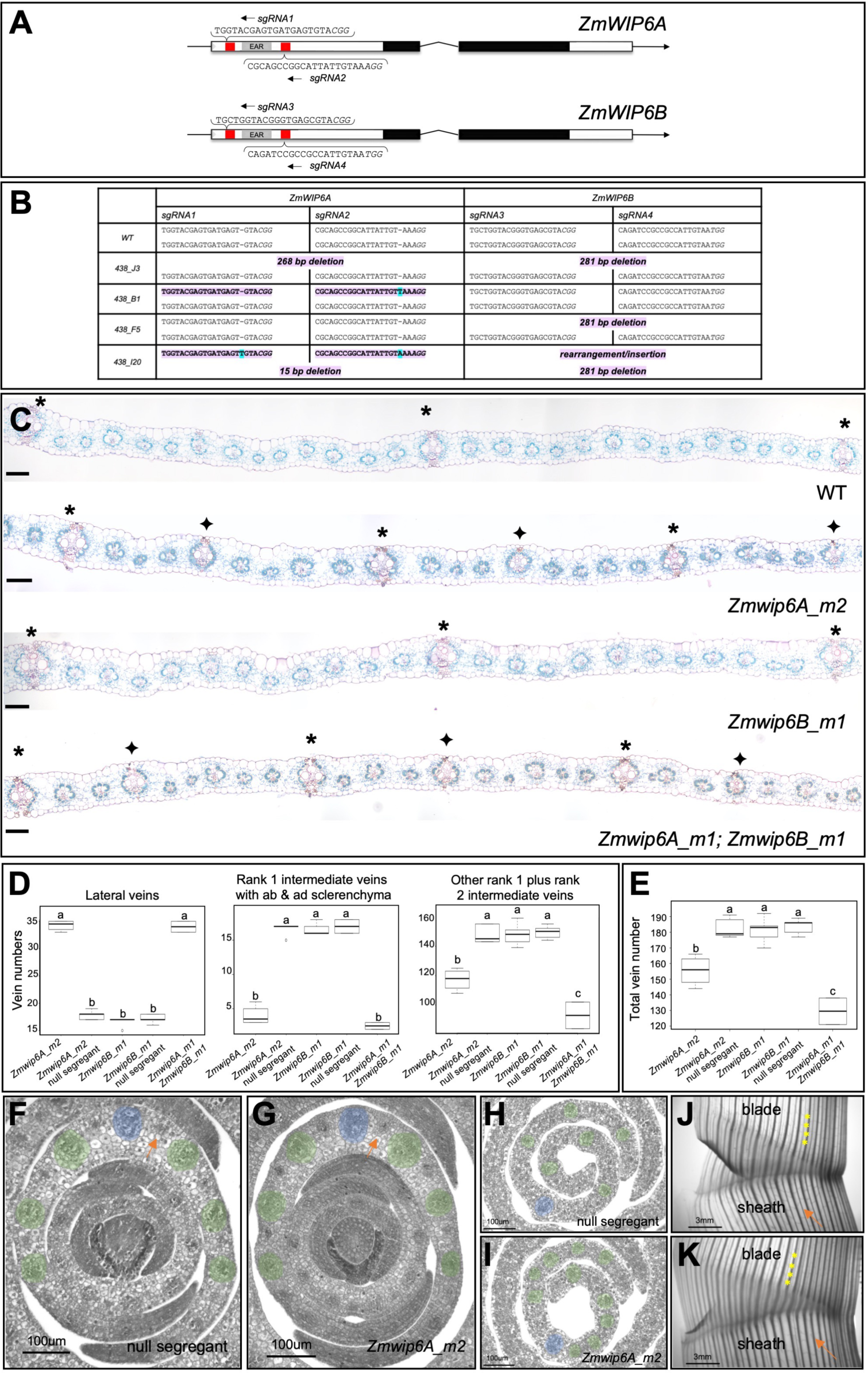
Loss of ZmWIP6A function leads to the formation of too many lateral veins in the maize leaf. **A)** Gene models of *ZmWIP6A* and *ZmWIP6B* showing 5’ and 3’ UTRs (black lines) plus two exons (rectangles) separated by an intron (^). The arrow indicates the direction of transcription, the conserved EAR domain (grey box), the C2H2 zinc finger region (black boxes) and the sgRNA target sites (red) are highlighted. The sgRNA sequences used for CRISPR/Cas9 mediated mutagenesis are as shown, with the PAM sequences in italics. **B)** Allelic composition of wild-type (WT) and four T1 maize lines. Mutant alleles are highlighted in pink and base insertions are highlighted in blue. One line (I20) was heteroallelic at both loci, one (J3) was heterozygous at both loci and the other two were heterozygous at one locus and wild-type at the other. **C)** Transverse sections of WT and mutant leaves showing vein numbers and types. In each case samples were taken from the 5^th^ leaf in the region between laterals 2 and 4, with lateral 1 being the closest to the midvein. Asterisks indicate lateral veins, diamonds indicate ectopic lateral veins. Scale bars = 100µm. **D, E)** Quantification of the number of each vein type (D) and of total vein number (E) across the entire medio-lateral axis of the 5^th^ leaf. ‘Other rank 1 veins’ are those which only have sclerenchyma on the adaxial or abaxial side but not both. N=5 for *Zmwip6B_m1*, *Zmwip6A_m2* null segregant and *Zmwip6B_m1* null segregant, N=4 for *Zmwip6A_m2* and N=2 for the double homozygous *Zmwip6A_m1/Zmwip6B_m1* mutant. Statistical significance was determined using a one way ANOVA followed by a Tukey’s test. **F-I)** Transverse sections of null segregant (F, H) and *Zmwip6A_m2* mutant (G, I) P4 primordia at the base (F, G) and in the middle (H, I) of the primordium. Midveins are highlighted in blue and lateral veins (both normal and ectopic) are highlighted in green. Orange arrows point to developing intermediate veins. Scale bars = 100µm. **J, K)** Cleared fragments of leaf 7 stained with Safranin showing the junction of leaf sheath and blade in null segregant (J) and *Zmwip6A_m2* (K) mutant leaves. Orange arrows point to a rank 1 intermediate with abaxial and adaxial sclerenchyma in (J) and an ectopic lateral in the equivalent position in (K). Yellow asterisks are positioned at transverse veins. Scale bar = 3mm.

To assess the phenotypic impact of loss of ZmWIP6 function, single and double mutant plants were grown alongside null segregants. None of the mutants exhibited general growth perturbations (**Figure S3C-F**), although *Zmwip6A_m2* mutants sometimes developed male inflorescences (tassels) with fewer fertile spikelets than segregating wild-type plants. In leaves, venation patterns were perturbed in single *Zmwip6A* mutants and in double mutants, but not in single *Zmwip6B* mutants (**Figure 3C**). Specifically, *Zmwip6A* and *Zmwip6A;Zmwip6B* mutant leaves developed twice as many lateral veins than corresponding wild-type siblings, with a concomitant decrease in the number of intermediate veins (predominantly rank 1 intermediates with sclerenchyma on both the adaxial and abaxial sides) (**Figure 3D**). Notably, the same phenotype was observed in leaves of *S. viridis* when the single *SvWIP6* gene was edited (**Figure S3C-F**). Given that the number of ectopic lateral veins was the same in single *Zmwip6A* mutants as in double mutants, no role could be inferred for *ZmWIP6B* in the regulation of vein type. When total vein number across the medio-lateral axis was quantified, however, *Zmwip6B* mutants were indistinguishable from wild-type but the two double mutants that were analyzed had fewer veins than single *Zmwip6A* mutants (**Figure 3E**). We thus conclude that *ZmWIP6B* plays a minor (as yet unidentified) role in venation patterning compared to *ZmWIP6A*. On the basis of the mutant phenotypes, we renamed the *ZmWIP6A* and *ZmWIP6B* genes *TOO MANY LATERALS* (*ZmTML1* & *ZmTML2*) and because of the conserved mutant phenotype, the *S. viridis* ortholog was named *SvTML*.

The development of extra lateral veins in *Zmtml1* mutants could result from the specification of additional lateral veins or from mis-specification of intermediate veins. To distinguish these possibilities, transverse sections of P4 primordia were examined at the base and middle of the primordium (**Figure 3F-I**). If the extra lateral veins result from ectopic specification, more laterals would be visible at the base of mutant P4 primordia than in null sgregants, because lateral veins differentiate from base to tip and are specified by P4. By contrast if the phenotype results from mis-specification of intermediate veins, developing veins sited in between lateral veins in the middle of mutant P4 primordia would be larger (more lateral-like) than in null segregants, because intermediate veins develop from tip to base and at P4 they are more differentiated in the middle than the base of the leaf. **Figures 3F, 3G** show that the number of lateral veins at the base of the P4 primordium is the same in null segregant and mutant samples. Developing veins positioned between the laterals look slightly larger in the mutant than in the null segregant but in both cases they are smaller than the adjacent lateral veins. Comparisons between **Figure 3H** and **3I**, however, reveal more veins with lateral identity in the middle of mutant primordia than in null segregants. Because ectopic lateral vein identity is evident in the middle but not the base of the primordium, we conclude that the mutant phenotype arises as a consequence of mis-specification of basipetally differentiating intermediate veins as lateral veins. This conclusion is validated because the rank 1 intermediate veins that differentiate from the tip of the blade into the sheath are clearly distinguishable from adjacent lateral veins in null segregants, but in mutant leaves they are not (**Figure 3J, K**). Notably, these longitudinal images also revealed no differences between mutants and null segregants in the pattern of transverse veins connecting lateral and intermediate veins across the medio-lateral leaf axis (**Figure 3J, K**). During wild-type development ZmTML1 must therefore act to suppress lateral vein identity in procambial centres that form rank 1 intermediate veins, specifically those rank 1 veins that are associated with sclerenchyma on both the adaxial and abaxial sides and extend from the leaf blade into the leaf sheath.

### Transcriptomes suggest altered cell division dynamics and auxin responses in *Zmtml1tml2* mutants

To identify potential targets of *ZmTML* function during early leaf development and to assess which processes are perturbed in mutant leaves, transcriptome profiles were compared between shoot apices (comprised of the shoot apical meristem and P1-P5 leaf primordia) of double *Zmtml1tml2* mutants and segregating wild-type siblings. Transcripts were quantified to identify those where levels showed a significant difference (log 2-fold change (L2FC) > 1; p < 0.05) between wild-type and mutant (**Supplementary File 8**). (All fold differences referred to below are L2FC at p < 0.05). Levels of transcripts encoded by 591 genes were significantly higher in mutant samples than wild-type and those encoded by 1275 genes were lower. GO term enrichment analysis of these 1866 genes revealed a broad range of functions that are perturbed in mutant plants, with those encoding transcription factors notably enriched for terms associated with shoot apical meristem development and adaxial/abaxial axis specification (**Figure 4A**). 89 transcripts representing 47 genes were up-regulated greater than 20-fold in the mutant, with the majority encoding enzymes or structural proteins (**Supplementary File 8**). By contrast, 45 transcripts representing 24 genes were down-regulated greater than 20-fold in the mutant and of the 11 genes in that group that have been annotated, 5 encode proteins that regulate gene activity (e.g. through ubiquitination, histone acetylation, histone methylation, chromatin remodelling or RNA silencing) (**Supplementary File 8**).

**Figure 4.**
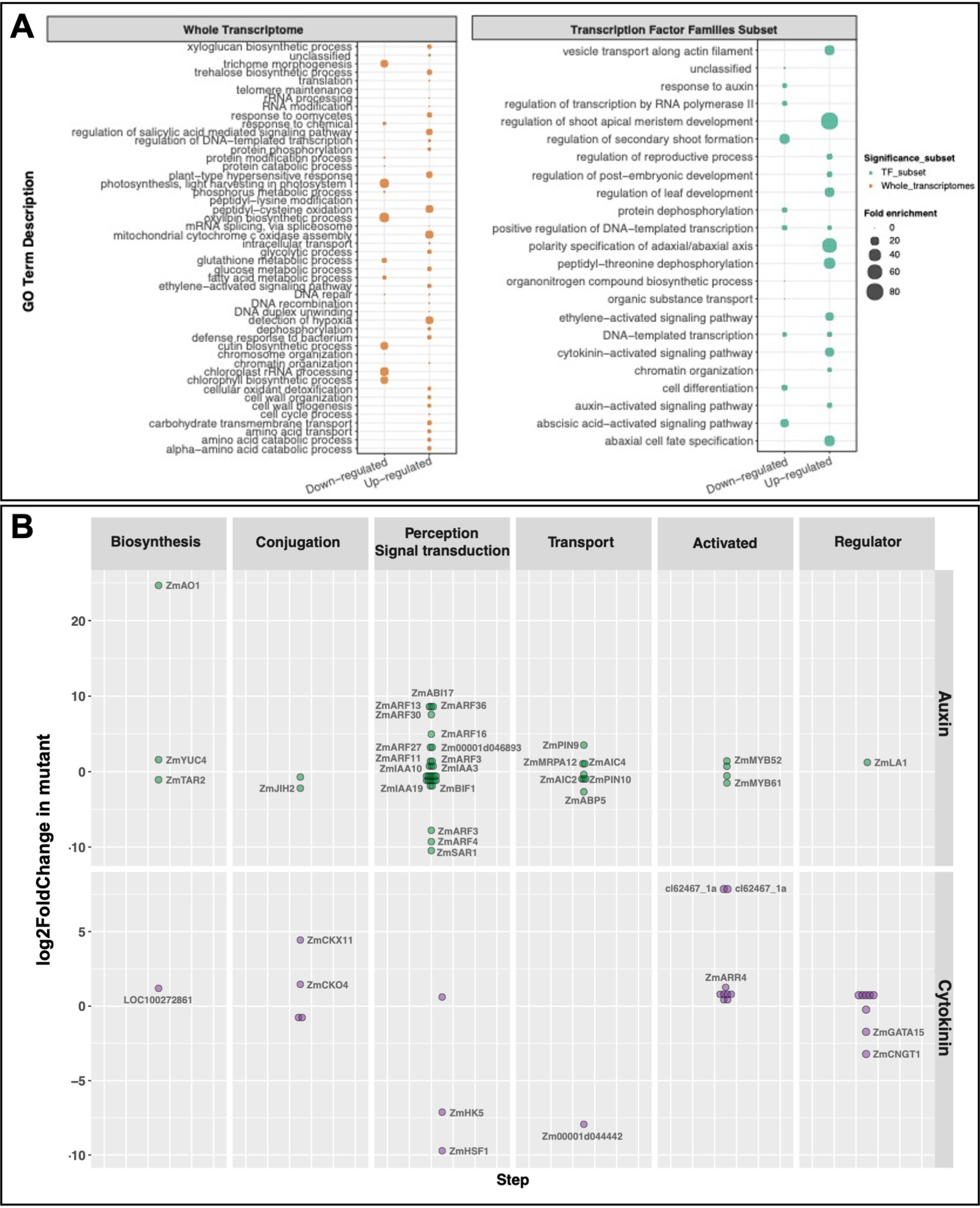
Evidence of auxin and cytokinin pathway perturbations in *Zmtml1tml2* mutant transcriptomes. **A)** Overrepresentation analysis (ORA) of significantly different transcript levels (L2FC >1; p ≤ 0.05) in *Zmtml1tml2* mutant apices as compared to corresponding segregating wild-type siblings, using data either from the whole transcriptome or from the subset annotated as transcription factors. Down- and up-regulated components are as indicated and the fold-enrichment in each case is shown by the size of the bubble. **B)** Log2fold difference in levels of transcript encoding auxin and cytokinin pathway components in *Zmtml1tml2* mutant apices as compared to corresponding segregating wild-type siblings. For each pathway step, the genes with significantly different transcript levels (L2FC ± 1; p ≤ 0.05) are labeled.

Of the few genes directly implicated in the development of Kranz anatomy (*SHORTROOT (SHR)* ^39^, *SCARECROW (SCR & SCR1h)* ^40–42^ and *NAKED ENDOSPERM (NKD1 & NKD2)* ^43^, only *NKD2* transcript levels were significantly different between mutant and wild-type (lower in the mutant - L2FC = 1; p < 0.05) (**Supplementary File 9**) suggesting that *ZmTML* genes act downstream of any ‘Kranz patterning module’. Notably, the failure to detect differences in levels of transcripts encoding orthologs of procambial markers in Arabidopsis (*ATHB8/ZmHB52* and *PIN1*) and rice (*OsHOX1/ZmHB78*) (**Supplementary File 9**) could also suggest that ZmTML function is downstream of procambium initial cell specification. The development of normal vein numbers in loss of function *Zmtml1tml2* mutants (**Figure 3D, E**) would also support this suggestion. However, *ZmHB52* is expressed more broadly than *ZmTML1* in wild-type leaf primordia, with transcripts detected across the P1 leaf primordium and in the more actively dividing cells at the margins of older primordia (**Figure S4A, S4B**). Restriction of *ZmHB52* transcripts to dividing procambial centres only occurs after *ZmTML1* transcript accumulation is repressed in those regions. We therefore propose that *ZmTML1* acts downstream of mechanisms that spatially regulate where veins are initiated, acts earlier than *ZmHB52* in vein development, and that any role in specifying procambial initial cells is carried out redundantly with other (as yet unidentified) partners such that procambium is positioned in the correct locations in mutant leaves.

Of the 529 differentially expressed transcription factor (TF) encoding genes (L2FC > 1; p < 0.05), greater than 4-fold differences in transcript levels between wild-type and mutant were observed for members of 24 different TF families (**Figure S4C**). Twenty-fold higher levels of *ZmGRF1* (*GROWTH REGULATING FACTOR 1*) transcripts in mutants than in wild-type segregants were particularly notable because *ZmGRF1* transcript levels are normally tightly regulated post-transcriptionally by miR396a ^44^. Whatsmore, constitutive overexpression of a miRNA-resistant version of *ZmGRF1* in maize was associated with a larger cell division zone at the base of developing leaves, with increased numbers of normal sized cells dividing more slowly than in wild-type counterparts ^45^. Upregulation of *ZmANT1* (*ZmEREB184*) (L2FC > 6; p < 0.05) in mutants (**Figure S4C**) is also notable in this regard because of a proposed role in cell proliferation during leaf development ^46^. As such, the phenotype of *Zmtml1tml2* mutant leaves may be caused in part by altered cell division dynamics in developing leaf primordia and *ZmGRF1* and/or *ZmANT1* expression may be directly or indirectly repressed by ZmTML1 and/or ZmTML2.

Because the spatial and temporal extent of cell division zones within developing grass leaves has been associated with the regulation of cytokinin signalling (CK) (higher levels are present in the division zone) ^47,48^, and GO-terms associated with CK were enriched in the population of differentially expressed genes (**Figure 4A**), we next sought to determine which components of the cytokinin pathway were perturbed in *Zmtml1tml2* mutants. To this end, we compiled a list of genes involved in cytokinin biosynthesis, signalling, degradation and response (**Supplementary File 10**), and quantified transcript levels for each in mutant and wild-type tissues. Of the 150 genes analysed, 11 exhibited significant differences between wild-type and mutant (**Supplementary File 10**, **Figure 4B**). The most substantial differences were seen for transcripts encoding two histidine kinase CK receptors (decreased in the mutant by more than 9-fold (*ZmHK1/HSF1* ^49^) and 7-fold (*ZmHK5*)), a homolog of the CK efflux transporter ABCG14 in Arabidopsis ^50^ (reduced more than 7-fold in the mutant) and the cytokinin oxidase degrading enzyme ZmCKX11 ^51^ (increased more than 4-fold in the mutant). Collectively these observations suggest that overall CK activity is repressed in *Zmtml1tml2* mutant apices and thus that cell division dynamics may be perturbed in the shoot apical meristem and/or leaf primordia.

Given that a role for auxin in leaf venation patterning is well established, we predicted that components of the auxin pathway would also be perturbed in *Zmtml1tml2* mutants. Indeed, significant differences were observed in levels of transcripts encoding a number of AUXIN RESPONSE FACTORs (ARFs) that bind directly to the promoters of auxin responsive genes (reviewed in ^52^) (**Figure S4C**). To determine the point(s) at which perturbations in the auxin pathway were manifest, we compiled a list of genes involved in auxin biosynthesis, conjugation, transport, signalling and response and quantified transcript levels for each gene (**Supplementary File 11**). Of the 186 genes analysed, 31 exhibited significant differences between mutant and wild type (**Figure 4B, Supplementary File 11)**. Although all steps of the pathway showed some perturbation, biosynthesis and transport steps were least impacted suggesting that these are upstream of TML function. With the exception of *ALDEHYDE OXIDASE1* (*ZmAO1*), where transcript levels were elevated over 20-fold in the mutant, the most substantial differences in transcript levels were seen for genes encoding components involved in auxin signalling. Aldehyde oxidases have been implicated in the synthesis of tryptophan precursors for IAA biosynthesis ^53^. However, direct functional analyses are lacking and given that there is no evidence that downstream components of the auxin biosynthesis pathway are similarly elevated we cannot speculate on the significance of any potential increase in tryptophan levels. Of the signalling components: *ZmSAR1*, *ZmARF4* and *ZmARF3* were downregulated more than 7-fold in the mutants; *ZmARF39/ZmABI17*, *ZmARF36* and *ZmARF13* were upregulated more than 7-fold in the mutant; and *ZmARF30* and *ZmARF16* were upregulated more than 4-fold in the mutant. Note that *ZmARF4* is a co-ortholog (with *ZmARF29*) of *MP* in Arabidopsis, mutations in which reduce the complexity of leaf venation patterns and disrupt axis formation in the embryo ^54,55^. *ZmSAR1* encodes the ortholog of Arabidopsis SUPPRESSOR OF AUXIN RESISTANCE 1, which is a nucleoporin that facilitates nuclear localization of AUX/IAA proteins ^56^. Reduced *ZmSAR1* transcript levels may thus suggest that the ability of AUX/IAA proteins to bind and repress ARFs is compromised in *Zmtml1tml2* mutants, which would translate into increased activation of ARF proteins. Notably, the three most highly upregulated *ARF* genes in *Zmtml1tml2* mutants are class B repressors and the two downregulated *ARF* genes are class A activators ^57^. The loss of ZmTML function may thus lead to an overall repression of auxin responses, but the possibility that responses are differentially up-or down-regulated in different spatial domains of the shoot apex cannot be discounted.

### Transgenic lines that constitutively express *ZmTML1* in rice could not be isolated

With the finding that *ZmTML1* represses lateral vein formation, we hypothesized that ectopic expression in rice would increase the number of intermediate veins in the leaf. Given that a higher proportion of intermediate veins is a key trait of C_4_ leaves, and an engineering goal is to make rice leaves more C_4_-like ^58,59^, we first tried to constitutively express *ZmTML1* in rice. Constructs in which expression of the maize *TML1* coding region was driven by the constitutive maize ubiquitin promoter (**Figure S5A**) were transformed on multiple occasions (n=7) and callus was propagated to the greening stage on selection medium, however, shoots could not be regenerated. This observation suggested that *ZmTML1* may suppress the formation of all veins and/or the pluripotency required for shoot regeneration.

### *TML* promoter activity is suppressed in transgenic rice lines

The failure to regenerate rice callus that constitutively expressed the *ZmTML1* gene from maize led us to consider that gene dosage and/or expression levels may need to be tightly regulated (spatially and/or temporally). To generate a chassis in which we could test this hypothesis, we therefore edited the endogenous rice gene. The rice genome contains a single *OsTML* (*OsWIP6*) gene (LOC_Os09g13680) which encodes a protein with 71.4 % amino acid identity to the maize protein. Edited alleles were generated either using a single guide RNA sequence (sgRNA1) that was designed to target between the EAR and zinc finger domains or by using sgRNA1 along with two additional guides (sgRNA3 and sgRNA4) that targeted the 5’ and 3’ regions of the coding sequence, respectively (**Figure 5A**). A total of five loss of function alleles were isolated from the two independent transformation experiments. The mutant alleles contained short indels at single or multiple guide sites that shifted the reading frame to result in aberrant or truncated proteins (**Figure 5B**). All edited T0 plants were either homozygous for the mutation or heteroallelic. After backcrossing to WT, heterozygous F1 plants were self-pollinated and F2 populations segregated mutant progeny as a recessive trait. Notably, all plants carrying loss of function alleles showed an increased proportion of larger, lateral-like veins in the leaf blade (**Figure 5C**) suggesting that *TML* gene function is conserved in maize and rice.

**Figure 5.**
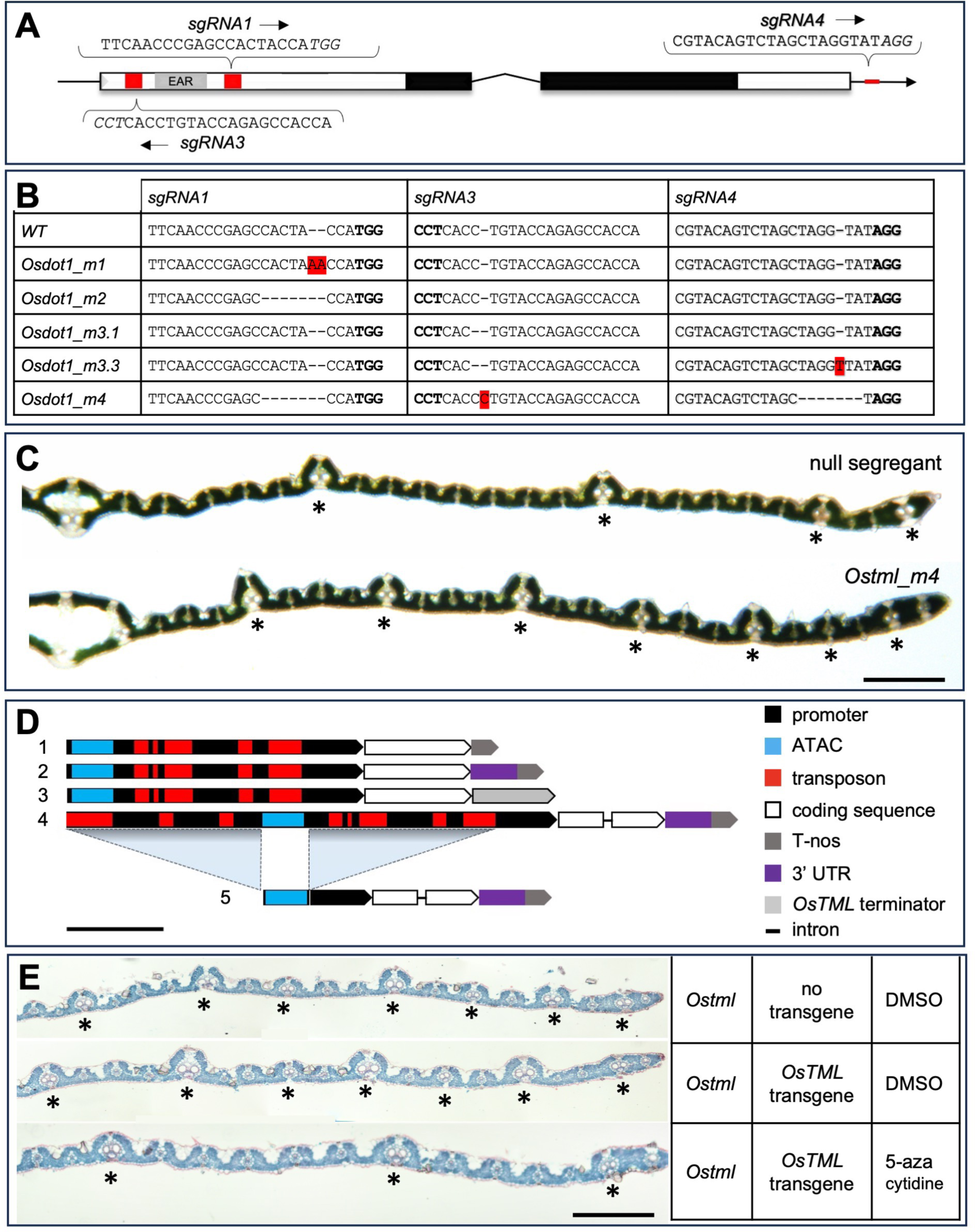
Ectopic lateral vein formation in *Ostml* mutants of rice cannot be complemented by *OsTML* because transgene expression is suppressed. **A)** Gene model of *OsTML* showing 5’ and 3’ UTRs (black lines) plus two exons (rectangles) separated by an intron (^). The arrow indicates the direction of transcription, the conserved EAR domain (grey box), the C2H2 zinc finger region (black boxes) and the sgRNA target sites (red) are highlighted. The sgRNA sequences used for CRISPR/Cas9 mediated mutagenesis are as shown, with the PAM sequences in italics. **B)** Allelic composition of wild-type (WT) and five mutant rice lines. Base insertions are highlighted in red. **C)** Transverse sections of null segregant and *Ostml_m4* mutant leaves (flag leaf minus 1). Asterisks indicate lateral veins. Scale bar = 700 µm. D) Schematics of 5 transgenes used to complement *Ostml* mutants. Scale bar = ∼1 kb. E) Transverse sections of leaf 7 from *Ostml_m3.1* mutant lines that were transformed with construct 2 from (D) and were germinated in the presence or absence of 50 mgL^-1^ 5 aza-cytidine. Asterisks indicate lateral veins. Scale bar = 300 µm.

The similar phenotypes of *Zmtml1* and *Ostml* mutants suggest that any TML-mediated difference in venation patterns between the two species must result from different domains of activity in the leaf. We thus revisited the question of whether maize-type expression patterns could alter venation patterns in rice and transformed the *ZmTML1* gene under the control of its own promoter (**Figure S5B, File S12**) into the *Ostml* mutant and wild-type Kitaake rice. Five hygromycin resistant transgenic lines were obtained in the *Ostml_m1* background and 24 lines in the wild-type background but no phenotypic changes were detected in any of the lines tested. This observation suggested either that the *ZmTML1* gene had recombined out of the transgene construct, that the *ZmTML1* promoter sequence did not contain essential regulatory regions, or that transgene expression was suppressed.

Given that we were unable to obtain transgenic lines that expressed *ZmTML* from either a constitutive promoter or from its own promoter, we tested whether endogenous *OsTML* expression patterns could be faithfully reproduced in a series of complementation experiments. First, a 2945 bp promoter sequence located directly upstream of the start codon and containing a regulatory region predicted by ATAC sequencing was fused to the *OsTML* coding sequence and to either the *nos* terminator (construct 1), the *nos* terminator plus 458 bp of the *OsTML* 3’UTR that contained an 83 bp element conserved in rice, maize and setaria (construct 2) or ∼1 kb of *OsTML* sequence downstream of the stop codon (construct 3) **(Figure 5D)**. The three constructs were individually transformed into wild-type and *Ostml_m1* (construct 1) or *Ostml_m2* and *Ostml_m3.1* (constructs 2 & 3) lines. In each case transgenic plants were successfully regenerated on selection medium but no complementation was observed (see examples in **Figure S5C, D**). As with *ZmTML1*, therefore, the transgenic rice promoter sequence was either lacking essential elements or expression was suppressed. To distinguish between these two possibilities, two further constructs were tested (**Figure 5D**). One contained the *OsTML* genomic region comprised of 5.35 kb of upstream sequence, both exons and the intron, plus 458 bp of downstream sequence (construct 4). Because the upstream genomic region contained multiple transposon sequences which may induce host silencing responses, these were removed in the second construct, leaving the ATAC sequence fused directly to the proximal promoter (construct 5). Both constructs were transformed into the *Ostml_m3.1* background but neither complemented the mutant phenotype (**Figure S5C, D**). Although still formally possible that essential elements were missing from the genomic constructs, this observation suggested that transgene expression was being suppressed, even in the absence of transposon sequences within the promoter. To test this hypothesis, seed from lines containing construct 2 were germinated on 50 mg/L 5-azacytidine to demethylate the genome. Although pleiotropic, this treatment lead to complementation of the *Ostml* mutant phenotype, presumably by activating *OsTML* transgene expression (**Figure 5E**). Collectively these observations suggest that *TML* gene expression is tightly regulated *in planta* and that any mis-expression (spatial and/or temporal) induced via transgenesis is likely to be suppressed or to cause developmental arrest.

### Loss of function mutations in *OsTML* specifically alter venation patterns in the leaf

Given the inability to modify *TML* expression patterns in rice, we characterized loss of function *Ostml* mutants in detail to determine the extent to which venation patterns are perturbed. Three independent lines were backcrossed to the wild type and selfed prior to examining overall growth phenotypes. In terms of overall morphology the mutant lines did not differ substantially from null segregants (**Figure 6A**) although *Ostml-m3.1* plants were shorter than wild-type (**Figure S6A**). All mutant lines showed a significantly increased number of lateral veins in the leaf blade (**Figure 6B, 6C**), a decreased number of intermediate veins (**Figure 6D**) but no significant change in vein density (**Figure 6E**). There were no apparent vascular defects in the roots, rachis or pedicel of mutant plants (**Figure S6B-D**) but venation defects in the leaf were apparent as early as P3, where veins developing in the position of intermediate veins were visibly larger in mutants than in corresponding wild types (**Figure S6E, F**). Importantly, the alterations to vein patterning were accompanied by altered bundle sheath cell occupancy in the leaf.

**Figure 6.**
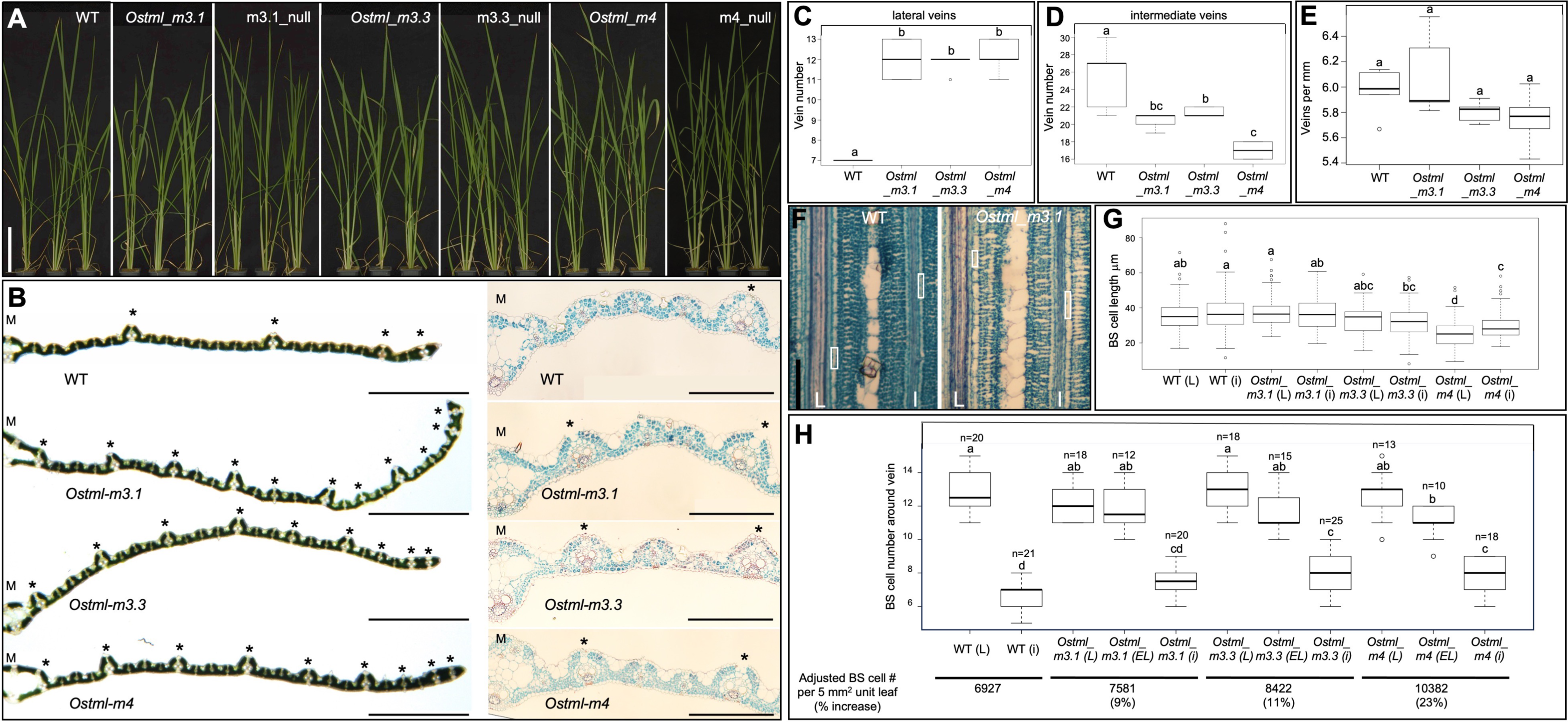
Venation defects in *Ostml* mutant leaves increase total vascular volume and alter bundle sheath cell dimensions. **A)** Whole plant phenotype of three independent *Ostml* mutant lines and respective null segregant siblings. Lines were selected from F2 populations that had been generated by backcrossing transgene free T1 mutants to Kitaake and then self-pollinating the resultant BCF1 progeny. Scale bar = 10 cm. **B)** Transverse sections imaged between the midvein and the leaf margin of flag leaf minus 1 (left panel) and from the midvein to the position corresponding to the first adjacent lateral vein in wild-type (WT) of leaf 4 (right panel). Asterisks indicate lateral veins. Scale bars = 1 mm (left panel); 200 µm (right panel). **C-E)** Quantification of lateral vein numbers (C), intermediate vein numbers (D) and vein density (E) in leaf 7 of WT and mutant plants. n = 5 for each genotype; different letters indicate statistically different groups as determined using a one-way ANOVA followed by a Tukey’s test (p value ≤ 0.05); see Supplementary File 13 for raw data. **F)** Paradermal images of the 7th leaf of WT and mutant plants, imaged in the middle of the blade. Lateral (L) and intermediate (i) veins are indicated. The longest bundle sheath cell around each vein in the field of view is outlined by a white box. Scale bar = 100 µm. **G)** Quantification of bundle sheath cell length around lateral (including ectopic laterals) and intermediate veins of WT and mutant leaves. n = 4 leaf samples and 82/145 cells around L/i veins for WT; 3 leaf samples and 96/114, 45/44 and 130/130 cells for *Ostml_m3.1, _m3,3* and *_m4* mutants respectively. Different letters indicate statistically different groups as determined using a one-way ANOVA followed by a Tukey’s test (p value ≤ 0.05); see Supplementary File 13 for raw data. **H)** Quantification of bundle sheath cells numbers around lateral (L), ectopic lateral (EL) and intermediate (i) veins in wild type and mutant plants. Different letters indicate statistically different groups as determined using a one-way ANOVA followed by a Tukey’s test (p value ≤ 0.05); see Supplementary File 13 for raw data. Median values for bundle sheath length, number of bundle sheath cells and total vein numbers per leaf were used to estimate total number of bundle sheath cells and percentage increase in number in a leaf unit of 5 mm wide and 1 mm long.

In wild-type rice, bundle sheath cells are rectangular when viewed paradermally, with the long edges running parallel to the associated vein and the short edges aligned with the medio-lateral leaf axis (**Figure 6F**). The length of individual bundle sheath cells around veins is highly variable but quantification revealed no significant differences between the length of cells around lateral versus intermediate veins in wild-type, *Ostml_m3.1* and *Ostml_m3.3* mutant samples (**Figure 6G**). The one exception to this was seen in *Ostml_m4* mutants where cells around both lateral and intermediate veins were significantly shorter than those in wild-type (**Figure 6G**). As a consquence of the increased proportion of lateral veins in mutant leaves, when viewed transversely, total bundle sheath cell number across the medio-lateral leaf axis is higher in all mutants as compared to wild-type (**Figure 6B, 6H**). When the altered cell number across the medio-lateral axis is adjusted to account for any variations in cell length along the proximo-distal axis, mutant leaves are seen to have an increase in bundle sheath cell number of 9-23% per unit area as compared to wild type (**Figure 6H**). Collectively these observations suggest that although there are no alterations in vein density in the *Ostml* mutant leaves, vein occupancy in the leaf is increased and there is a substantial increase in the number of bundle sheath cells.

## DISCUSSION

Through loss of function analyses in three monocot species we have identified a conserved role for WIP6 orthologs in the specification of vein rank in developing grass leaf primordia **(Figures 3**, 5, 6 **& S3)**. WIP6 orthologs have been identified in the genomes of all vascular plant groups, as have members of the sister WIP1-5 clade, but just a single *WIP* gene is present in the non-vascular bryophyte *Marchantia polymorpha* ^60^ (**Figure 1**). The finding that the *MpWIP* gene is required for the development of air pore complexes on the dorsal surface of the thallus led to the suggestion that prior to a gene duplication in the last common ancestor of lycophytes and euphyllophytes (ferns, gymnosperms and angiosperms), WIP gene function was required for the differentiation of multicellular structures ^60^. This suggestion is consistent with the observation that members of the WIP1-5 clade in Arabidopsis have been shown to play a role in the development of transmitting tract tissue in the pistil (WIP2) ^61^, the seed coat (WIP1) ^31^, and the root (WIP2/4/5) ^62^, specifically in the regulation of cell division orientation in stem cells and cell-type specification. However, a role for WIP6 has not been unambiguously defined in Arabidopsis. Original reports of leaf venation defects in loss of function mutants ^32^ were later invalidated ^34^ and the suggestion of a non cell-autonomous role in maternal tissue to suppress cell-type specification in embryonic roots was only inferred from the phenotype of combined triple *wip1/3/6* mutants ^33^. To our knowledge, there are no reports of phenotypic perturbations in single *wip6* loss of function mutants in any eudicot.

On the basis of the conserved phenotype observed in loss of function *wip6* mutants in maize, *S. viridis* and rice, we named WIP6 orthologs *TOO MANY LATERALS* (*TML*). The spatial expression of *TML* genes in procambial initial cells, the temporal restriction of expression to the first few divisions of those cells and the mis-specification of intermediate veins as laterals in mutant leaves (**Figures 2, 3, 5, 6, S2, S3 & S4**) are all consistent with a role in cell-type specification and/or divisions of procambial cells in the leaf. The mutant phenotype is also consistent with the role inferred from triple *wip1/3/6* mutants in Arabidopsis, namely that of a suppressor ^33^. A repressor role is further supported by the observation that constitutive expression of *AtWIP6* conditions a small leaf phenotype in Arabidopsis ^63^. Notably, general repression of organ growth by WIP proteins has also been demonstrated in other species. For example, in melon the WIP1 ortholog *CmWIP1* regulates sex determination by repressing carpel primordia in male flowers ^64^ and overexpression of the WIP2 ortholog in *Gerbera hybrida* suppresses growth in multiple organs ^65^. This general repressive role likely explains the failure to regenerate rice callus after transformation with a construct that constitutively expressed *ZmTML1* (**Figure S5**). It may also explain why attempts to express *TML* genes in rice from the corresponding maize or rice promoters led to transgene silencing **(Figure 5)**. Possibly, the repressive function of TML is only tolerated in the first few divisions of procambial cells in leaf primordia and as such any leaky expression in callus is silenced. Considering all of these factors, we propose that TML functions in grasses to suppress cell divisions and/or cell fate decisions in procambial cells that normally lead to the formation of lateral veins, consequently enabling intermediate veins to develop.

The role of TML as a suppressor of lateral vein formation as opposed to an activator of intermediate vein formation is intriguing, not least because it only functions in a subset of the rank 1 intermediate veins. All intermediate veins differentiate from the tip of the leaf primordium towards the base, but only the largest and first to initiate extend into the leaf sheath ^17,18^. It is these large intermediate veins that are converted to laterals in *tml* mutants of both maize and rice. The rank 1 intermediate veins that only differentate sclerenchyma on the abaxial or adaxial side and the rank 2 intemediate veins that only develop in C_4_ leaves are unaffected in mutant leaves **(Figures** 3, 5 & 6). This phenotypic distinction in mutants is reflected in the spatial distribution of *TML* transcripts during wild-type development **(Figures 2 & S4)** and thus regulators that activate *TML* transcription and or transcript stability determine which procambial cells are subject to TML-mediated suppression. To date only one transcription factor that binds *in vitro* to the *ZmTML1* promoter has been identified ^66^ and no function has been assigned to *ZmbHLH105* or any of its orthologs. Although direct activators of TML activity are unknown, at some level the PIN1/auxin traces that precede the formation of all vein ranks ^24^ must be interpreted differently in procambial initial cells that develop the different vein-types, leading to TML activity in some but not all of those cells.

Transcriptome comparisons between wild-type and mutant shoot apices in maize have provided some insight into perturbations caused by loss of TML function, however, because *TML* is expressed in just a few cells and only for a short time, the likelihood of identifying direct targets in this dataset is low. Observed perturbations in both auxin and cytokinin pathways **(Figures 4 & 4S)** could be a direct consequence of loss of TML function but there is insufficient spatial resolution in whole tissue transcriptomes to validate this suggestion. Specifically, the apparent suppression of both pathways observed in whole shoot apices could be masking significant increases in activity that are spatially or temporally localized. Mechanistically, the presence of an EAR domain ^38^ in the TML protein is of note because EAR domains bind TOPLESS proteins, which themselves recruit epigenetic modifiers ^37^. It is therefore possible that promoters bound by the zinc finger domain of TML are silenced. In this regard, a 20-fold increase in levels of transcripts encoding the growth regulating factor *ZmGRF1* in *Zmtml1tml2* mutant leaves is notable. Although direct targets have yet to be identified, we propose that TML either represses a target gene that positively regulates lateral vein identity or that TML-mediated regulation of cell divisions in procambial initial cells prevents specific division orientations that are required for lateral vein formation.

Unlike in eudicots, where venation patterning in the leaf has been perturbed by mutations in a number of genes that are not directly associated with auxin signaling (e.g. *cotyledon vascular pattern* ^67^ and *thickvein* ^68^) most venation defects in grass leaves reported to date have resulted from indirect perturbations to hormone pathways, mis-specification of blade versus sheath domains, and/or the regulation of leaf width. For example, growth of maize on auxin transport inhibitors converted blade tissue into sheath, with a consequent alteration in vein spacing across the medio-lateral leaf axis ^69^. A similar blade to sheath transition was observed in sorghum mutants defective in a cytochrome P450 required for brassinosteroid biosynthesis ^70^ and in gain of function rice mutants that lead to an up-regulation in cytokinin signaling ^71^. In rice, mutants with higher vein density invariably had narrower leaves and smaller mesophyll cells between veins ^72,73^ and in the case of the *narrow leaf 1* mutation, that defect was associated with reduced polar auxin transport ^74^. The only mutation reported to directly perturb procambium formation in rice leaves (*radicleless1*), also failed to form an embryonic root ^75^. Notably, this pleiotropic phenotype is also observed in the Arabidopsis auxin response mutant *monopteros* but *MP* is known to play a direct role in the leaf vein formation ^55^. In contrast to all of these examples, the role of *TML* in vein development is confined to the leaf, providing an opportunity to manipulate vein patterning in rice for future metabolic engineering of C_4_ photosynthesis. An optimal chassis for C_4_ engineering would have normal leaf width, vein pairs separated by just two mesophyll cells and a 1:1 ratio of bundle sheath to mesophyll cells. Loss of TML function does not achieve that goal, however, mutant leaves have a significant increase in the overall volume of leaf veins and in the number of bundle sheath cells. The leaf-specific defects in *Ostml* mutants thus represent a fundamental advance in our understanding of how veins are patterned in grass leaves and provide the first step towards engineering the anatomy required to underpin C_4_ photosynthesis.

## METHODS

### Phylogenetic analysis

For the WIP family tree, the complete set of predicted proteomes for all species in Phytozome version 12 ^76^ were subject to orthogroup inference using OrthoFinder ^30^. The orthogroup containing the maize gene GRMZM2G150011(v3 genome)/ Zm00001d020037(v4 genome)/Zm00001eb309530 (v5 genome) was identified and the 379 constituent protein sequences (**Supplementary File 1**) subject to multiple sequence alignment using MergeAlign ^77^. The alignment was then imported into MEGA ^78^ and trimmed to contain the EAR domain plus the zinc finger region. After empirical testing of the multiple sequence alignment for maximum likelihood phylogenetic tree inference using IQTREE ^79^, 11 sequences that failed the composition χ^2^ test and 21 sequences that had more than 50% gaps were removed. After realignment of the remaining 347 sequences using MergeAlign (**Supplementary File 2**), the best-fitting model parameters (JTT+I+G4) were estimated and a consensus phylogenetic tree was estimated from 1000 bootstrap replicates (**Supplementary File 3**). The data were imported into ITOL ^80^ to generate the pictorial representation. Branches with less than 50% bootstrap support were deleted and three leaves that were incorrectly positioned in the WIP2/4/5 clade (likely due to mis-annotation of the corresponding sequences) were deleted (Asparagus v1 05.3078 – positioned in a eudicot clade, *H. annus* chr02g0056311 – positioned in a monocot clade, and *Zostera marina* Zosma112g00240.1 – splitting gymnosperm and Amborella).

For the WIP6 tree, sequences were selected from two bryophytes (to root the tree), 6 eudicots and 7 monocots (**Supplementary File 4**). Species were chosen primarily to resolve relationships within the Panicoid grasses but also to represent the major flowering plant groups (monocots, asterids and rosids). Sequences were retrieved by BLAST searches of Phytozome 13 and NCBI and aligned with MAFFT using the L-INS-i refinement method ^81^ (**Supplementary File 5, Supplementary** Figure 1A). IQTree ^79^ was used to estimate the best-fitting model parameters (JTT+F+I+G4) and to infer a consensus phylogenetic tree from 1000 bootstrap replicates (**Supplementary File 6**). The data were imported into ITOL to generate the pictorial representation. Notably it was extremely difficult to align sequences between the EAR and WIP domains across a broad phylogenetic range, with blocks of sequence conserved within groups (e.g. grasses and brassicas) but highly variant between groups (**Supplementary** Figure 1B**, Supplementary File 7**). Despite a high level of sequence conservation across the zinc finger domains, this variability constrained the number and identity of eudicot species for which tree inference was supported with high bootstrap values.

### Plant material and growth conditions

Maize (inbreds B73 and B104) and *Setaria viridis* (accession ME034V) seeds were germinated as previously described ^41,43^. Both species were grown in a greenhouse with a long day (16 h light / 8 h dark) photoperiod with 28 °C day / 20 °C night temperatures. Under low natural light conditions (below 120 µmol photon m^-2^ s^-1^) plants were provided with supplemental light. Rice (*spp japonica cv* Kitaake) seeds were sterilized and germinated in petri dishes containing ½ MS medium under a 16 h light /8 h dark photoperiod, 30 °C day/ 25 °C night temperatures. Seven days after plating, seedlings were transferred to falcon tubes containing ¼ MS medium. Two-week-old seedlings were transferred to clay pots and grown under the same conditions described above.

*In situ hybridization In situ* hybridization was carried out using wax-embedded shoot apices as described by Schuler et al. ^82^, with digoxygenin (DIG)-labelled RNA probes designed to specifically detect *ZmWIP6A, ZmWIP6B, OsWIP6, SbWIP6* or *SvWIP6* transcripts. The *ZmWIP6A* probe was a 427 bp fragment spanning 141 bp of the 5’UTR and the first 286 bp of the coding sequence. The probe shared 53% similarity with the corresponding region of the *ZmWIP6B* transcript sequence. The *ZmWIP6B* probe was a 203 bp fragment encompassing the end of the coding sequence and part of the 3’UTR. The probe shared 52% similarity with the corresponding region of the *ZmWIP6A* transcript sequence. The *OsWIP6* (142 bp), *SbWIP6* (393 bp) and *SvWIP6* (214 bp) probes encompassed the beginning of the first exon and included the EAR domain. Post-hybridization washes were undertaken with 0.005 x SSC buffer made from a 20x SSC stock (3M NaCl, 0.3M Na_3_citrate), calculated to ensure stringency.

### Molecular Cartography probe design

Probes were designed using Resolve BioSciences’ proprietary design algorithm and gene annotations from the Zea Mays RefGen V4. The final set of probes was selected with the methodology described in Glenn et al. ^83^.

### Tissue preparation and Molecular Cartography hybridization

Wild-type B73 seeds were germinated and grown in an environment-controlled chamber on a 16 h/8 h day/night cycle, temperature 28 °C/20 °C, 50 % humidity and light intensity of 300 µmol m^-2^ s^-1^. Seeds were germinated in medium vermiculite and seedlings were harvested ten days after sowing. Maize shoot apices were cut just below and 0.5 cm above the shoot apical meristem and immediately fixed in 4 % paraformaldehyde, dehydrated, and embedded in Paraplast Plus following the user guide for sample preparation from Resolve BioSciences. Clearing and wax infiltration were either carried out manually or in a TissueTek VIP processor. Wax-infiltrated samples were embedded in blocks of Paraplast Plus using a TissueTek TEC wax-embedding centre. A total of six 10 µm transverse sections across two experiments were cut with a microtome and baked onto a Molecular Cartography-compatible coverslip at 37 °C overnight to allow for sample attachment. Sections were then deparaffinized, permeabilized, and refixed according to the user guide. After complete dehydration, the sections were mounted using SlowFade-Diamond Antifade mounting medium, covered with a glass coverslip and shipped to Resolve BioSciences for analysis.

At Resolve BioSciences, sections were washed twice in phosphate buffered saline for 2 min, followed by 1 min washing in 50% and 70% ethanol at room temperature. Ethanol was removed by aspiration and DST1 buffer was added, followed by tissue priming for 30 min at 37 °C and by a 48 h hybridization using probes specific for the target genes. Samples were washed to remove excess probes and fluorescently tagged in a two-step color development process. Fluorescent signals were removed after imaging in a decolorization step. Colorization, imaging, and decolorization were iterated for multiple cycles to generate a unique combinatorial code for each target gene. Samples were imaged by Resolve BioSciences as described in Glenn *et al.* ^83^.

### Pre-processing of Molecular Cartography images and cell segmentation

Cell segmentation was carried out using CellPose 2.0 ^84^, starting from high resolution images of Calcofluor and DAPI-stained sections. A pre-processing step was carried out to facilitate cell segmentation. Firstly, DAPI and Calcofluor images were overlaid in ImageJ: the co-existence of cell wall and nuclear staining allows for more accurate segmentation with CellPose. Subsequently, different stages of leaf development were cropped and isolated from the overlay image: this allows for faster computational speed during the segmentation step, and for a more accurate estimate of the average cell diameter, which varies greatly between leaf primordia. The images of isolated leaf primordia were imported into CellPose, and the Calcofluor and DAPI channels brightness and contrast were adjusted as required. The “cyto2” model was used to predict the cell boundaries in meristem/P1 and P2 primordia, whereas the “TN2” model was used to predict cell boundaries in P3, P4 and P5 primordia. All segmentations were then manually corrected, and the resulting masks were used to train a custom model. All subsequent primordia underwent segmentation using the custom models thus obtained and corrected manually where required. Segmented cells were exported as PNG masks and converted to ROIs using the ImageJ plugin “Labels to ROIs”. ROIs were then used to visualize single-cell gene expression using the proprietary Molecular Cartography plugin from Resolve Biosciences.

### CRISPR construct design

Short RNA guides targeting *ZmWIP6A, ZmWIP6B, SvWIP6 and OsWIP6* were designed using the online guide RNA selection tool, CRISPOR ^85^. For maize and setaria, two guide RNAs were designed to target each *WIP6* gene within the first exon. In the case of *OsWIP6* three guides were used, sgRNA1 and sgRNA3 in the first exon and sgRNA4 in the 3’UTR to potentially generate large deletions in combination with the first two guides.

The four maize guides targeting *ZmWIP6A* (sgRNA25: TGGTACGAGTGATGAGTGTA & sgRNA194: CGCAGCCGGCATTATTGTAA) and *ZmWIP6B* (sgRNA28: TGCTGGTACGGGTGAGCGTA & sgRNA309: CAGATCCGCCGCCATTGTAA) were cloned in vector pMHb-pZmUBIL-ZmCas9-NosT-AG-pBdEF1a-tdTomato-NLS-NosT (https://gatewayvectors.vib.be/).

The rice and setaria sgRNAs were cloned downstream of the OsU3 promoter in the Golden Gate system ^86^ as previously described ^42^. The setaria guides sgRNA77: CTCATCCTACTCGGCATGCT and sgRNA153: CCGTTCCCCCCAGCAACAAG were assembled in construct EC17847. The rice guides were used to generate two separate constructs, EC17627 containing only sgRNA1: TTCAACCCGAGCCACTACCA and EC17697 containing sgRNA1, sgRNA3: CACCTGTACCAGAGCCACCA and sgRNA4: CGTACAGTCTAGCTAGGTAT. The Cas9p gene ^87^ was expressed under the control of the *ZmUbi* promoter and the selection module contained a hygromycin resistance gene driven by the *OsAct1* promoter.

### Overexpression and complementation construct design

The *ZmUbi_pro_*: *ZmTML1* overexpression construct was obtained by cloning the *ZmTML1* ORF into the binary destination vector pVec8-Gateway ^88^. The *ZmTML1* ORF was amplified by polymerase chain reaction (PCR) using maize B73 genomic DNA as template and Gateway® compatible primers (MS. F: 5’-TCAGCAGACCACCACCCAAT-3’; MS. R: 5’-CTAGATGACGACCATGTCGCTGG-3’). The PCR product was cloned into Gateway® donor vector pDONR™207, sequenced, and subsequently cloned downstream of the *ZmUBI* promoter in the binary destination vector pVec8-Gateway.

*OsTML* complementation constructs 1, 2 and 3 were cloned into the Golden Gate system which required BpiI or BsaI enzyme recognition sites to be domesticated. The *OsTML* cDNA sequence was obtained by synthesis with one BpiI site ‘domesticated’. The 2945 bp promoter region, *OsTML* terminator region and the 3’ UTR were cloned from PCR amplified fragments. As the *OsTML* promoter contained two BsaI and one BpiI enzyme recognition sites, primers were designed both to ‘domesticate’ the restriction sites (by introducing three single bp changes) and to generate BpiI sites at the ends. This allowed promoter assembly using a Golden Gate cloning strategy and without introducing any other scars in the sequence. The *OsTML* terminator region and 3’UTR did not require domestication.

To avoid any modification that could potentially alter promoter function, the fourth version of the complementation construct was assembled using NEBuilder HiFi DNA Assembly strategy. A ∼7 kb genomic region was amplified by PCR in four overlapping fragments. The fragments were assembled into a Golden Gate level 2 vector containing the hygromycin selection module. To obtain the ‘ATAC’ version (construct 5), the construct above was cut with PmeI/FspI to remove most of the promoter sequence apart from 545 bp adjacent to the ATG. A PCR amplified 540 bp fragment overlapping the predicted ‘ATAC’ region was then added in a NEBuilder HiFi DNA Assembly reaction. The ATAC sequence information for both maize ^89^ and rice ^90^ was accessed via the Plant Epigenome Browser - https://epigenome.genetics.uga.edu/PlantEpigenome/index.html. Primer sequences for Golden Gate and NEBuilder HiFi DNA Assembly are listed in **Supplementary Table 1.**

### Plant transformation

Maize CRISPR lines were generated in inbred line B104 at the VIB Ghent Crop Genome Engineering Facility (https://www.psb.ugent.be/cores/crop_genome_engineering_facility). Backcrossed T1 seeds were provided for 10 independent lines.

Transgenic rice (*O. sativa ssp. japonica* cv. Kitaake) lines were obtained following a modified callus transformation protocol from Toki et al. ^91^. The modified protocol is available at https://langdalelab.files.wordpress.com/2015/07/kitaake_transformation_2015.pdf.

To generate setaria CRISPR edited lines, construct EC17847 was transformed into accession ME034V following a protocol described by Hughes *et al.* ^43^.

Regenerated rice and setaria T0 plantlets were verified by PCR with primers targeting the selection gene hygromycin phosphotransferase (hpt) (forward primer: 5’-CAACCAAGCTCTGATAGAGT-3’; reverse primer: 5’-GAAGAATCTCGTGCTTTCA-3’) to confirm successful transgene insertion in the genome. Plants genotyped positive for hpt were potted in clay and watered with fertilizer solution in a semi-hydroponic setting.

### Phenotyping leaf anatomy

Leaf segments (3-4mm) spanning the whole leaf width were collected from the widest part of fully expanded leaves, which in maize and setaria localized to the middle of the blade and in rice to the upper third of the blade. Leaf seven was analysed in rice, and leaf five in maize and setaria. The samples were fixed in 3:1 ethanol: acetic acid for 30 min at room temperature and transferred to 70 % ethanol. Fixed samples were placed into tissue embedding cassettes, dehydrated and infiltrated with Paraplast Plus in a Tissue Tek Vacuum Infiltration Processor. Wax infiltrated samples were set in blocks, trimmed and sectioned using a Leica RM2135 rotary microtome. 10 μm transverse sections were placed onto microscope slides and dried at 37 °C overnight before dewaxing and staining. For paradermal sections, fragments of rice leaf blade without the midvein were flattened and stacked using 0.8 % agarose layers before wax embedding and sectioning as described above.

Leaf sections were treated with histoclear to remove wax, partially rehydrated in 70 % EtOH prior to staining for 30 min with Safranin O (1 % in 50 % EtOH). The slides were then stained for 30 sec with Fast Green FCF (0.03 % in 95 % EtOH), dehydrated, drained and mounted using DPX.

Sections were viewed and imaged with a Leica DMRB microscope. Images were used to estimate leaf width, vein density and quantification of each vein type (lateral, rank1 intermediates and rank 2 intermediates). Paradermal sections were imaged to allow bundle sheath cell length to be measured.

### RNA sequencing, transcript quantification and differential expression

Maize shoot apices were dissected from vermiculite-germinated 7-day-old *Zmtml1tml2* seedlings and genotypically segregating wild-type siblings. Total RNA was extracted from three individual shoot apices using the Qiagen RNeasy Plant Mini Kit (Cat. No. 74904). Subsequent DNase treatment was done using the Ambion® TURBO DNA-free™ Kit (Cat. No. AM1907). RNA quantity was checked and normalized using a Qubit™ RNA HS Assay Kit. RNA quality was assessed with an Agilent RNA 6000 Nano Kit ensuring that all RNA used for library preparation had a RIN value ≥ 9.0. Strand-specific RNA sequencing library preparation and sequencing was conducted by the Oxford Genomics Centre at the Wellcome Centre for Human Genetics (funded by Wellcome Trust grant reference 203141/A/16/Z). Sequencing was performed on an Illumina NovSeq6000 with paired-end 150-bp reads. Sequenced reads were mapped to the Zea Mays B73 RefGen V4 genome (https://phytozome-next.jgi.doe.gov/info/Zmays_RefGen_V4) using STAR ^92^, transcript quantification was done with Salmon ^93^ and differential gene expression was determined using DESEq2 ^94^. The identification of differentially expressed genes was based on having log 2-fold change (L2FC) ±1, with an adjusted p-value <0.05.

### Differential expression data sub-setting and enrichment analysis

A subset of genes that were differentially expressed between *Zmtml1tml2* plants and genotypically segregating wild-type siblings were classified as transcription factors using the Grassius Maize Transcription Factor Database (https://grassius.org/browsefamily/Maize/TF) annotations. Lists of B73 RefGen V4 IDs of genes belonging to each specific transcription factor family were downloaded and cross-referenced with the differentially expressed gene list using R scripts (available upon request). Genes known to have a role in auxin and cytokinin pathways were compiled and curated from searches of both the literature ^95^ and databases (AmiGo knowledgebase ^96^, PANTHER ^97^ and UniProt ^98^) using the search words “auxin” or “cytokinin”.

### Enrichment analysis

Statistical overrepresentation test (ORA) of Gene Ontology annotation set “GO biological process complete” was performed using the PANTHER 17.0 GO Ontology database version (DOI: 10.5281/zenodo.7942786 Released 2023-05-10) with a Fisher’s Exact test type and a false discovery rate correction. The reference list comprised of all significantly differentially expressed genes between the *Zmtml1tml2* plants and genotypically segregating wild-type siblings. The significantly up-regulated (L2FC > 1; padj <0.05) and significantly down-regulated (L2FC < 1; padj <0.05) datasets were tested separately. A subset of the differentially expressed genes between *Zmtml1tml2* plants and genotypically segregating wild-type siblings were classified as transcription factors using the Grassius Maize Transcription Factor Database ^99^ annotations. An independent ORA was conducted using this subset with the reference list comprising all significantly differentially expressed genes between the *Zmtml1tml2* plants and segregating wild-type siblings. Enriched parent GO terms that were overrepresented at FDR <0.005 and a corrected P value of ≤ 0.05 were plotted in R using ggplot2 ^100^.

### Statistical Analysis

Statistical tests were undertaken using R Studio. One-way ANOVA tests were performed for analyses involving more than two groups, followed by Tukey’s HSD post-hoc testing if the ANOVA *p* value was <0.05. Details of ANOVA analyses are summarized in Supplementary File 13.

### Data access

All read datasets were deposited at the NCBI sequence read archive (http://www.ncbi.nlm.nih.gov/sra) under accession numbers SRSxxxxx–SRSxxxxx inclusive.

## LIST OF SUPPLEMENTARY MATERIAL

**Figure S1.** WIP6 protein alignment.

**Figure S2.** Spatial localization of WIP6 transcripts in different grass species.

**Figure S3.** WIP6 loss of function alleles in maize and setaria.

**Figure S4.** Transcription factor expression profiles in maize shoots.

**Figure S5.** Transgenic rice lines expressing *ZmTML* or *OsTML* could not be generated.

**Figure S6.** Venation defects are restricted to leaves in *Ostml* mutants.

**Table S1.** Primer sequences used for cloning *OsTML* complementation constructs.

**File S1.** List of sequences in the WIP orthogroup.

**File S2.** Alignment of 379 WIP protein sequences used to infer the phylogeny in Figure 1A.

**File S3.** IQTreefile of WIP phylogeny presented in Figure 1A.

**File S4.** Subset of 17 WIP6 protein sequences used to infer the phylogeny in Figure 1B.

**File S5.** Alignment of 17 WIP6 protein sequences (illustrated in Figure S1A) and used to infer the phylogeny in Figure 1B.

**File S6.** IQTreefile of WIP6 phylogeny presented in Figure 1B.

**File S7.** Subset of 50 WIP6 protein sequences from 41 species used for the alignment in Figure S1B.

**File S8.** List of genes that are differentially regulated in shoot apices of wild-type and *Zmtml1tml2* mutants.

**File S9.** Levels of transcripts encoding candidate regulators of Kranz anatomy in wild-type and *Zmtml1tml2* mutant shoot apices.

**File S10.** Genes encoding cytokinin pathway components that are differentially regulated in wild-type and *Zmtml1tml2* mutant shoot apices.

**File S11.** Genes encoding auxin pathway associated genes that are differentially regulated in wild-type and *Zmtml1tml2* mutant shoot apices.

File S12. Alignment of *ZmTML1* promoter with the domesticated version used in transformation constructs.

**File S13**. Data and statistical analyses supporting main figures.

## AUTHOR CONTRIBUTIONS

D.V., C.P., M.Z., O.S. and J.A.L. conceived and designed the experiments; J.A.L. carried out the phylogenetic analysis (Figs. 1 & S1), O.S. performed *in situ* hybridizations (Figs. 2 & S2); C.P. and S.B. generated the molecular cartography data (Figs. 2, S2 & S4), M.Z. performed the transcriptome analysis (Figs. 4 & S4), D.V. carried out all of the remaining experiments. D.V., C.P., M.Z. and J.A.L. analysed the data. D.V. and J.A.L. wrote the first draft of the manuscript and all authors contributed to the final version.

## ACKNOWLEDGEMENTS

The authors thank Laurens Pauwels and the VIB team for maize transformation; David Emms for providing the WIP orthogroup sequences; Mara Schuler for making the *ZmUBI_pro_:ZmTML* construct; John Baker for plant photography; Roxaana Clayton, Julie Bull, Lizzie Jamison and Nina Johnson for technical support; Tom Hughes, Sophie Johnson, Tina Schreier, Sovanna Tan and Steve Kelly for discussion throughout the experimental work and during manuscript preparation.

---

**Supplementary Figure 1.**
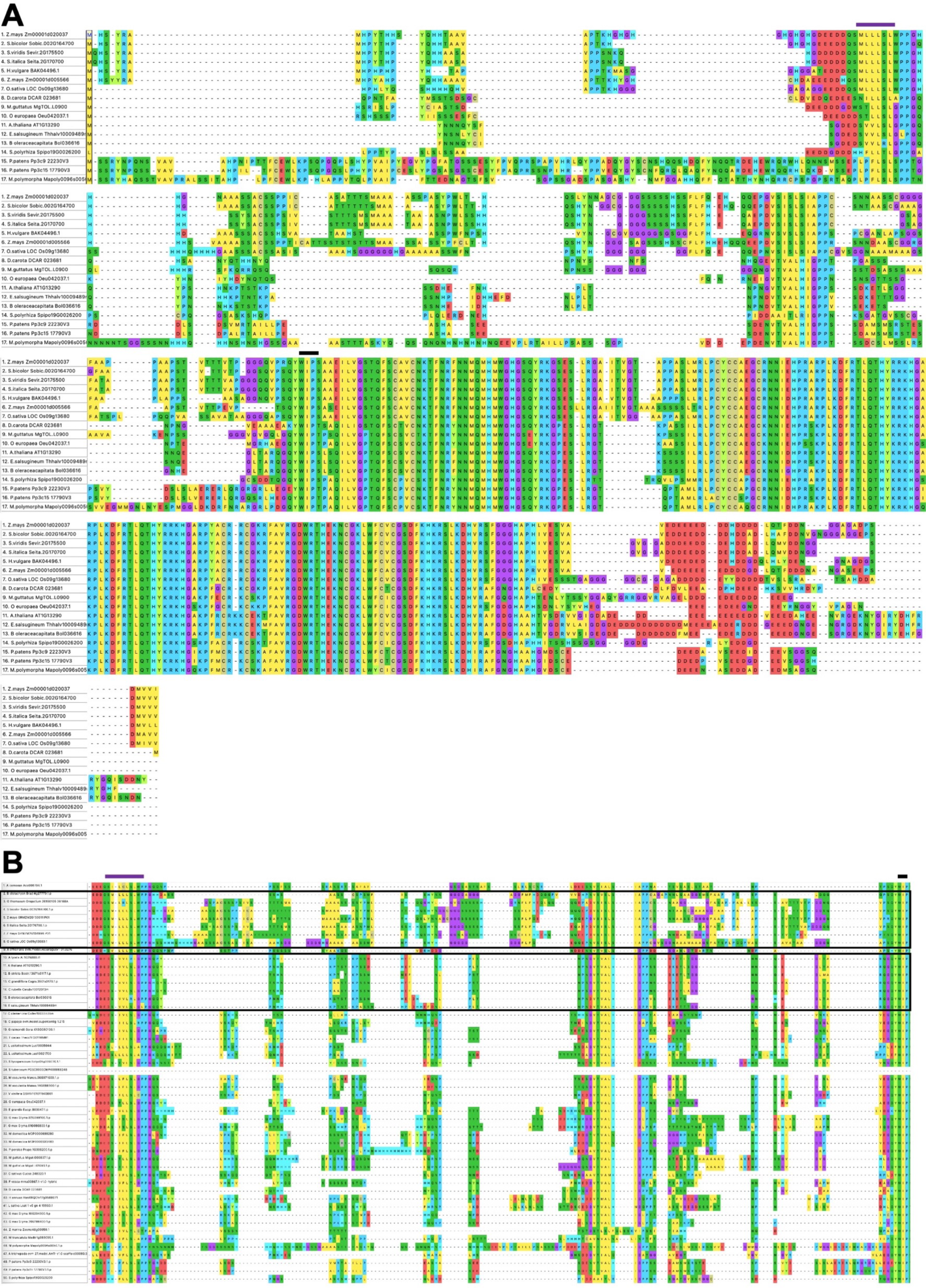
WIP6 protein alignment. **A, B)** Alignment of WIP6 sequences used to infer phylogenetic tree in Figure 1B (A) and to represent a broader range of species (B). Horizontal lines ove the alignment denote the EAR (purple) and WIP (black) domains. Note that sequences in between EAR and WIP domains do not align across a broad phylogenetic range, although blocks of sequence conserved within groups e.g. grasses (top box in (B)) and brassicas (bottom box in (B)).

**Supplementary Figure 2.**
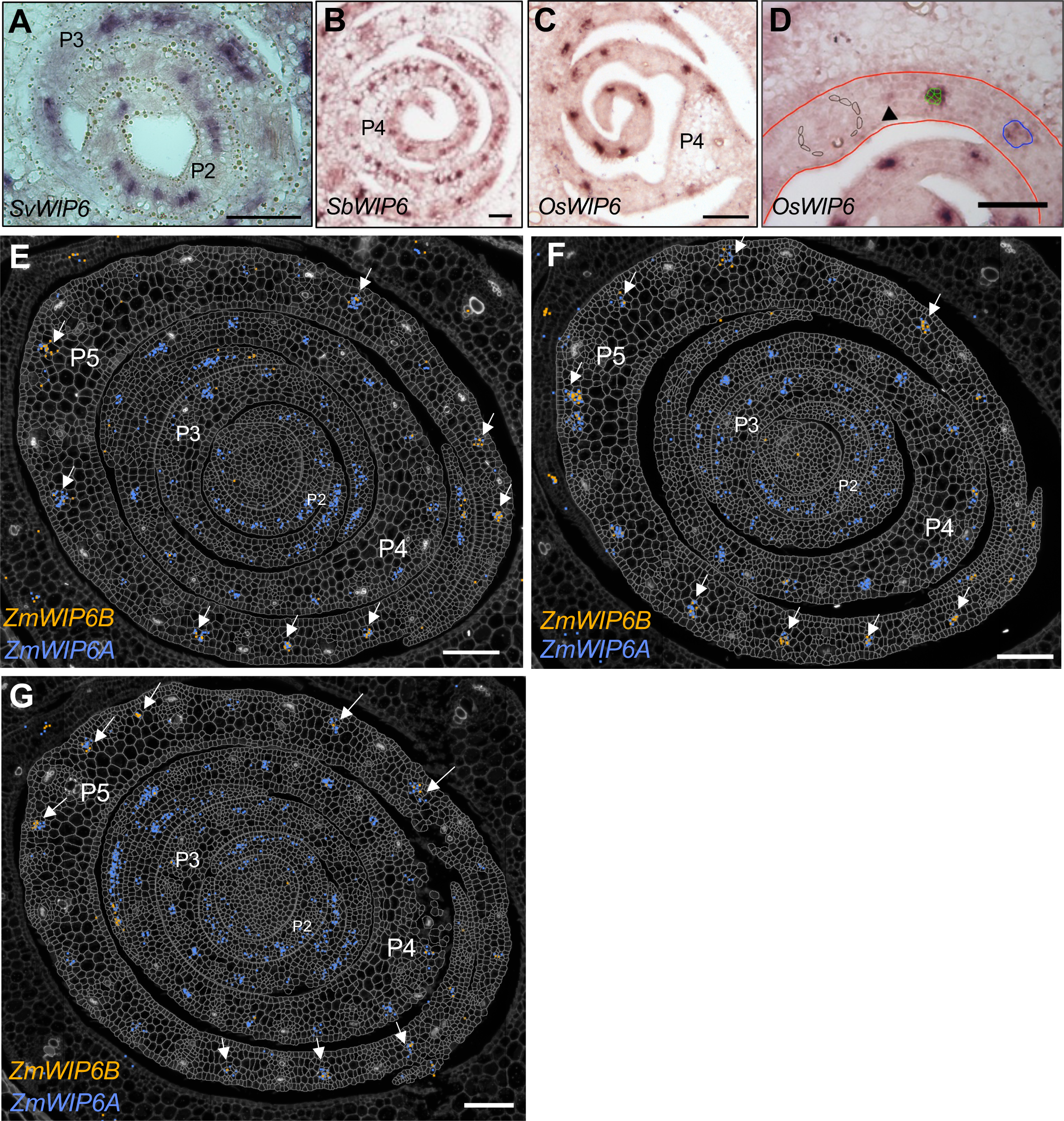
Spatial localization of WIP6 transcripts in different grass species. **A-C)** *In situ* hybridization to shoot apices of *Setaria viridis* (A), *Sorghum bicolor* (B) and *Oryza sativa* (C). Plastochron stages are as indicated. Scale bars = 50 μm. **D)** Annotated image of a section hybridized with *OsWIP6*. The P4 primordium is outlined in red. Four stages of vein development are highlighted. Arrowhead marks a procambial initial; green cells an early dividing procambial centre; blue line outlines a more advanced stage and black cells surround a fully differentiated lateral vein. *OsWIP6* transcripts are detected in the procambial initial and at high levels in the early dividing procambium. Lower levels are evident later in vein development and transcripts are absent from fully differentiated veins. Scale bar = 50 μm **E-G)** Molecular cartography of *ZmWIP6A* (blue) & *ZmWIP6B* (orange) transcripts in P1-P5 leaf primordia of maize. White arrows indicate developing intermediate veins in the P5 leaf sheath tissue. Scale bar = 100μm.

**Supplementary Figure 3.**
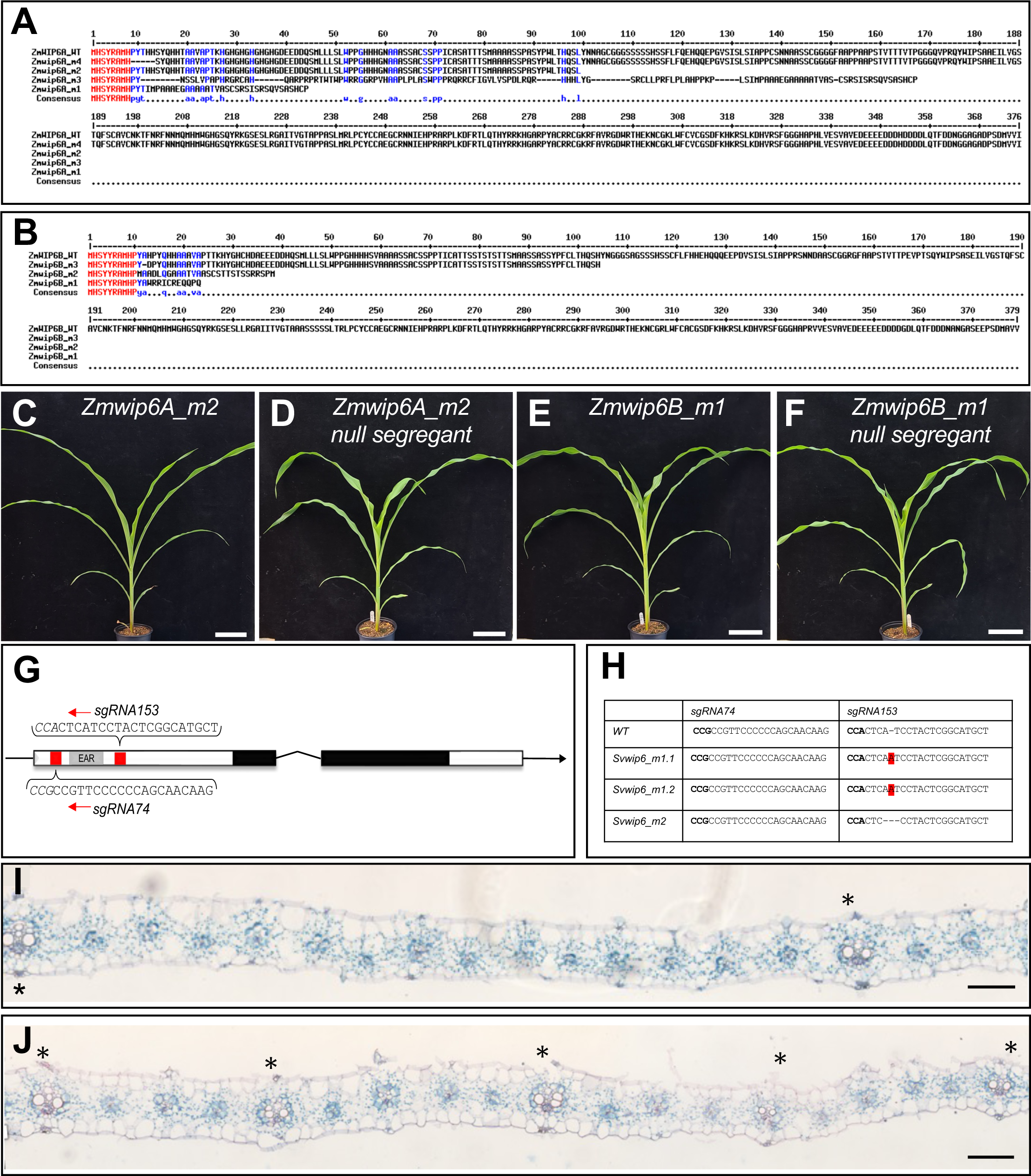
WIP6 loss of function alleles in maize and setaria. **A, B)** Amino acid sequence alignments showing predicted proteins encoded by *Zmwip6A* (A) and *Zmwip6B* (B) mutant alleles. **C-F)** Phenotype of ∼one month old *Zmwip6A* mutant (C) and null segregant (D), and *Zmwip6B* (E) mutant and ll segregant (F) plants. Scale bar = 10 cm **G)** Gene model of *SvWIP6* showing showing 5’ and 3’ UTRs lack lines) plus two exons (rectangles) separated by an intron (^). The arrow indicates the direction of nscription, the conserved EAR domain (grey box), the C2H2 zinc finger region (black boxes) and the RNA target sites (red) are highlighted. The gRNA sequences used for CRISPR/Cas9 mediated mutagenesis are indicated with the PAM sequence in italics. **H)** Summary of wild-type (WT) and *Svwip6* utant alleles. **I, J)** Transverse sections of wild-type *S.viridis* and *Svwip6* mutant leaves. Asterisks indicate lateral veins. Scale bar = 100 mm.

**Supplementary Figure 4.**
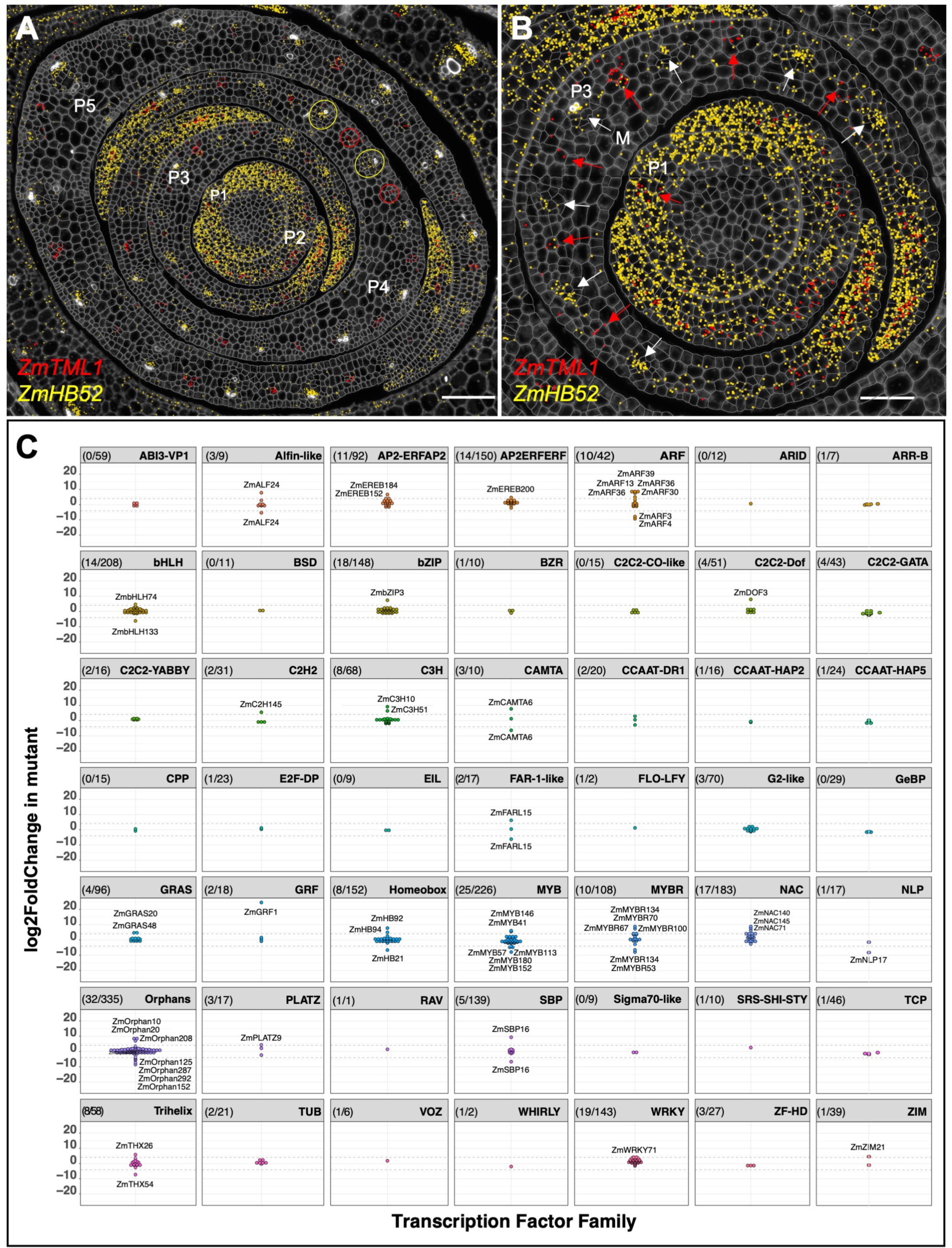
Transcription factor expression profiles in maize shoots. **A, B)** Molecular cartography of *ZmTML1* (red dots) and *ZmHB52* (yellow dots) transcripts in the shoot apical meristem and P1-P5 leaf primordia of wild-type maize apices. Yellow and red circles indicate discrete regions of *ZmHB52* and *ZmTML1* transcript accumulation in differentiated lateral (yellow) and developing intermediate (red) veins of P4 leaf sheath. White arrows indicate foci of *ZmHB52* transcript accumulation in the P3 primordium at the midvein (M) and at positions where developing procambial centres comprise ∼6 cells. Red arrows indicate *ZmTML1* transcript accumulation in the P3 primordium at sites of newly initiated procambial centres that comprise ∼1-4 cells. At P1, *ZmHB52* transcripts are detected throughout the primordium whereas *ZmTML1* transcripts are restricted to the developing midvein (red arrow). Scale bars = 100μm (A); 50μm (B). Section is same as in Figure S2A. **C)** Log2fold difference in levels of transcripts encoding transcription factors in *Zmtml1tml2* mutant apices as compared to corresponding segregating wild-type siblings. For each transcription factor family, the number of genes with significantly different transcript levels (L2FC **±** 1; p ≤ 0.05) between mutant and wild-type is indicated relative to the number of genes in the family in parentheses. Gene names are indicated where more than 4-fold differences were detected.

**Supplementary Figure 5.**
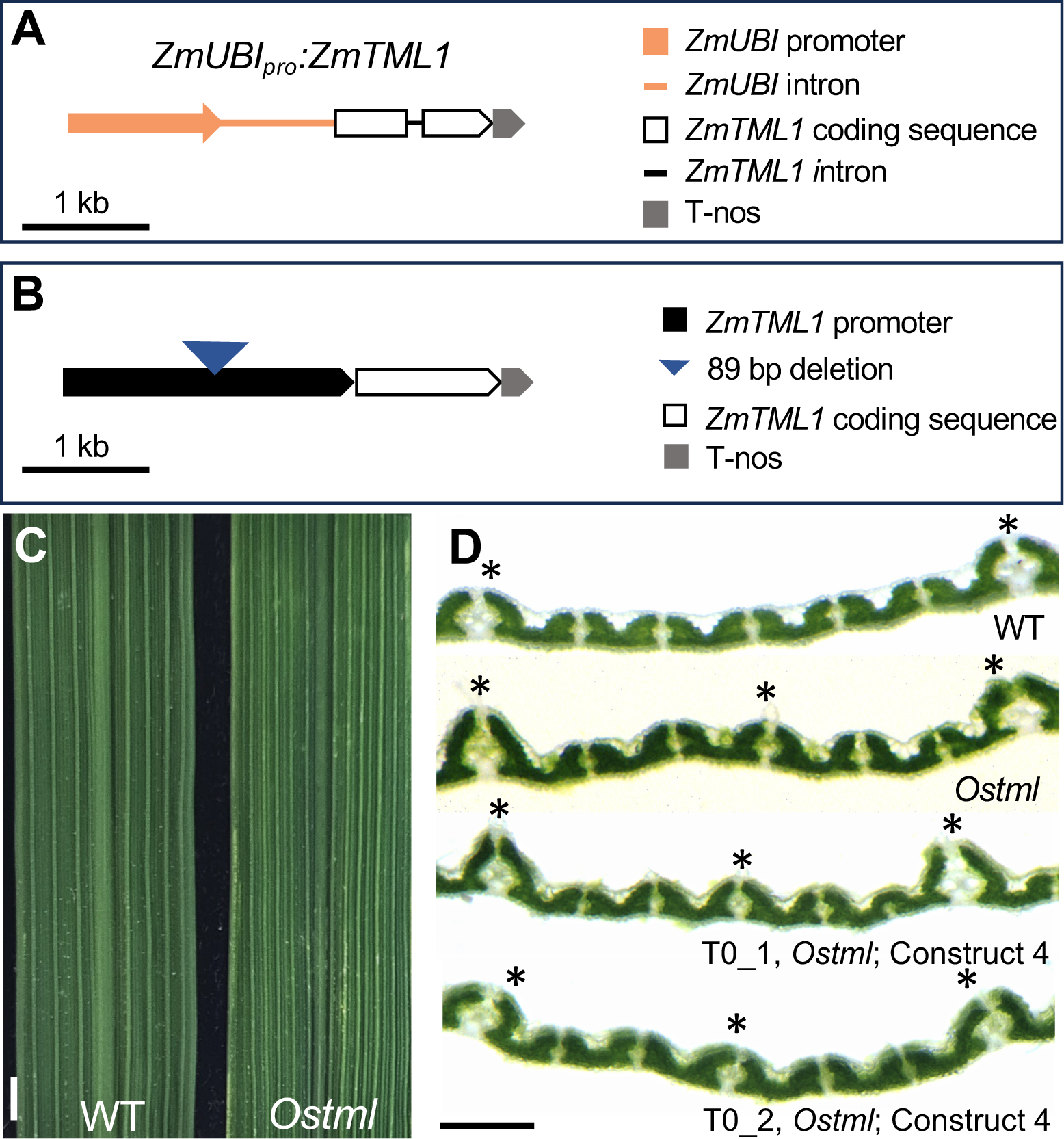
Transgenic rice lines expressing *ZmTML* or *OsTML* could not be generated. **A)** Schematic of construct in which expression of the *ZmTML1* coding sequence was driven constitutively by the maize ubiquitin (*UBI*) promoter. **B)** Schematic of construct used to test *ZmTML1* promoter in rice. **C)** Flag leaf phenotype of wild type and *Ostml* plants showing differences in vein patterns. Scale bar = 2 mm. **D)** Free-hand transverse sections of leaf 7 used to screen transgenic lines in complementation experiments, in this case transformed with complementation construct 4. No complementation was observed in the absence of 5-aza cytidine treatment. Scale bar = 100 µM.

**Supplementary Figure 6.**
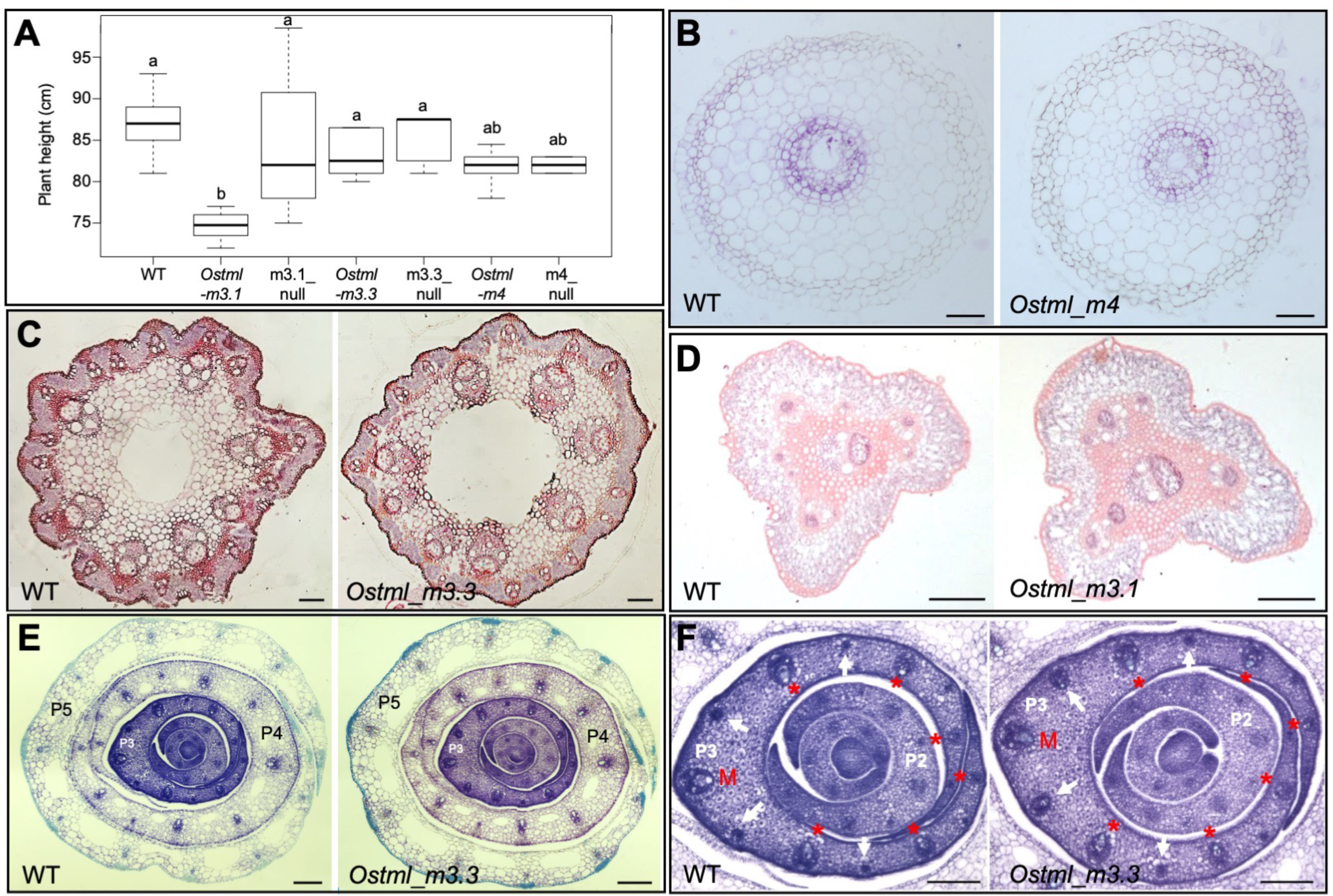
Venation defects are restricted to leaves in *Ostml* mutants. **A)** Quantification of plant height. n ≥ 4; different letters indicate statistically different groups as determined using a one-way ANOVA followed by a Tukey’s test (p value ≤ 0.05); see Supplementary File 12 for raw data. **B-D)** Vein patterning is unperturbed in roots (B), rachis (C) and pedicel (D) of *Ostml* mutants. **E, F)** Transverse sections of shoot apices showing the shoot apical meristem and P1-P5 primordia. Venation patterns are indistinguishable in wild-type (WT) and mutant leaf sheaths (visible for P4 and P5 in (E)). In developing leaf blades of P2 and P3 primordia, the size and position of the midrib (M) and lateral veins d asterisks) are normal but the veins developing at the position of intermediate veins (white arrows) are larger in the mutant (F). Scale bars = 100 µm.

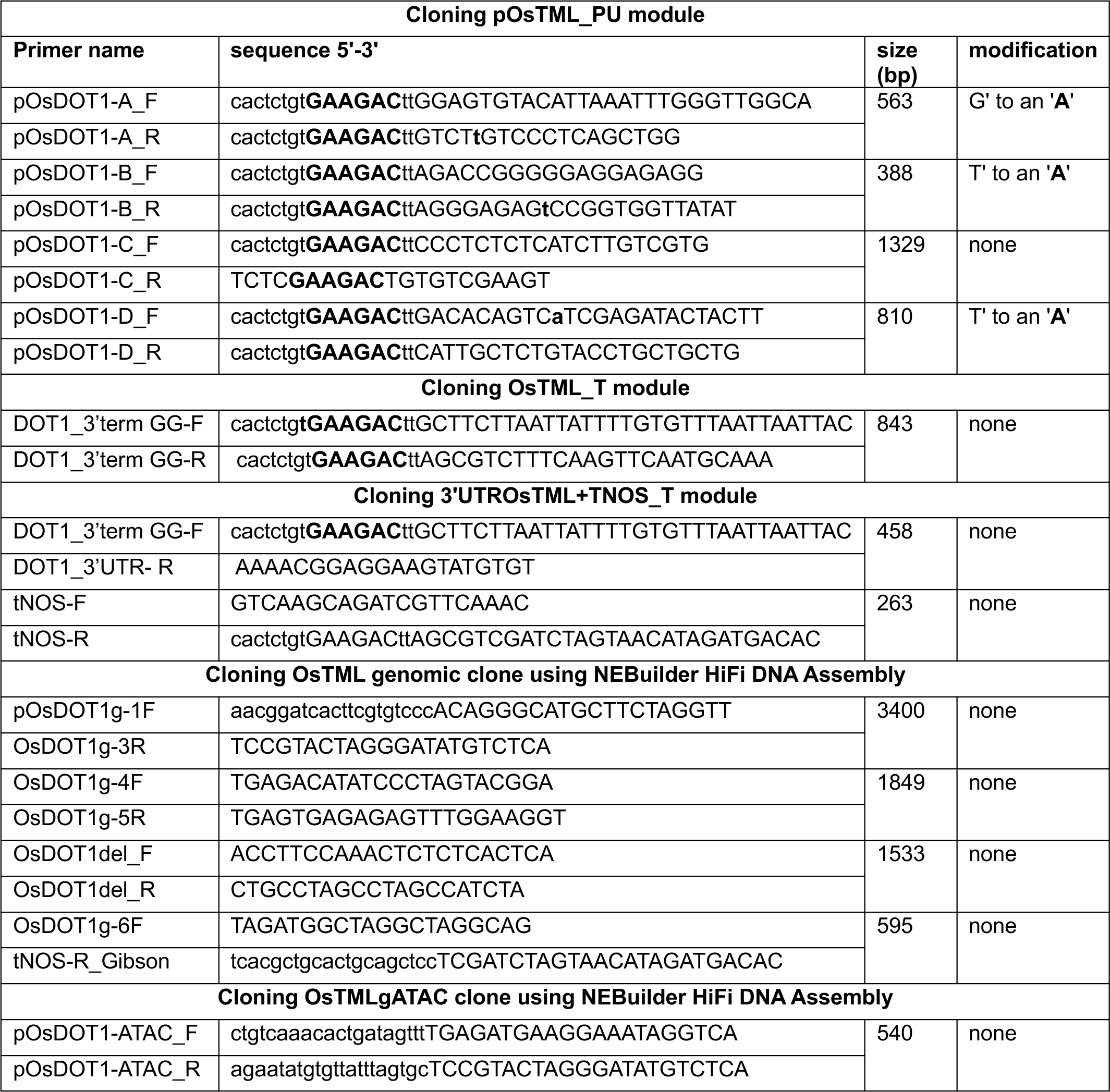
Supplementary Table 1. Primer sequences used for cloning the *OsTML* complementation constructs. Domestication of the BsaI/BpiI enzyme recognition sites was done by introducing base pair changes listed under “modification”. The BpiI sites used for assembling the fragments into a Level 0 OsTML PU module are marked in bold. Primers for the NEBuilder assembly carry 20 bp primer extensions (lower case) to allow seamless integration into the destination vector.

