## Supplementary material for "THE WIP6 TRANSCRIPTION FACTOR *TOO MANY LATERALS* SPECIFIES VEIN TYPE IN C_4_ AND C_3_ GRASS LEAVES": File S12

**Supplementary File 12. Alignment of *ZmTML1* promoter with the domesticated version used in transformation constructs**

```
Clustal alignment
2500bp_upstream      CGACTAATTTGTTCCGACGGGCTAACTTCCGACGGTTTCGGCTAACTTCCGACGGTTTAA	60
GG_promoter          CGACTAATTTGTTCCGACGGGCTAACTTCCGACGGTTTCGGCTAACTTCCGACGGTTTAA	60
                     ************************************************************

2500bp_upstream      ATAACTTCCGAGAGTTTTGCGCTAACTTCCTACGGTTTAGGCCCTCAGAAGTTAAGTATT	120
GG_promoter          ATAACTTCCGAGAGTTTTGCGCTAACTTCCTACGGTTTAGGCCCTCAGAAGTTAAGTATT	120
                     ************************************************************

2500bp_upstream      TTGGTGTAGTGTGTTATATTATAGTTCTCTGAATTTCAAGGCAAGAATTGTATATAACTT	180
GG_promoter          TTGGTGTAGTGTGTTATATTATAGTTCTCTGAATTTCAAGGCAAGAATTGTATATAACTT	180
                     ************************************************************

2500bp_upstream      CCTAGCGGTACAATCAAAAGAAACGATTCACATTTTTTTCTATAGTCAAAATAATGGCAA	240
GG_promoter          CCTAGCGGTACAATCAAAAGAAACGATTCACATTTTTTTCTATAGTCAAAATAATGGCAA	240
                     ************************************************************

2500bp_upstream      ATTACATGTTGCCAATGAGGCACTTTATAGGAGAAGTGCCTAAGGCTATGGCCAGCAGAT	300
GG_promoter          ATTACATGTTGCCAATGAGGCACTTTATAGGAGAAGTGCCTAAGGCTATGGCCAGCAGAT	300
                     ************************************************************

2500bp_upstream      CATACATATGGTCGTGCAAAACACTGTTTACAGAGTATAATTTGAAATGAGAGATATATG	360
GG_promoter          CATACATATGGTCGTGCAAAACACTGTTTACAGAGTATAATTTGAAATGAGAGATATATG	360
                     ************************************************************

2500bp_upstream      AAGATGAGTAACCTGCTAGAGATAGCCTTAAGTGAACTAAATGCTCATCCAAAAAAATTG	420
GG_promoter          AAGATGAGTAACCTGCTAGAGATAGCCTTAAGTGAACTAAATGCTCATCCAAAAAAATTG	420
                     ************************************************************

2500bp_upstream      TGTGAAACCTATATTTATGATTGCTTCTGGAAGACTTTGTTTTTATTATTGCTTTAAAAG	480
GG_promoter          TGTGAAACCTATATTTATGATTGCTTCTGGATGACTTTGTTTTTATTATTGCTTTAAAAG	480
                     ******************************* ****************************

2500bp_upstream      AAAGATTTTATTTCATTCCAAGATTAGTACATCGAGTTGATACATGGTCTTAGAATAGTC	540
GG_promoter          AAAGATTTTATTTCATTCCAAGATTAGTACATCGAGTTGATACATGGTCTTAGAATAGTC	540
                     ************************************************************

2500bp_upstream      TCCCAGCAGCCTCTTCTATGTGATAGACACACAACTTAAGATTTAAAATATTCTAAGCAT	600
GG_promoter          TCCCAGCAGCCTCTTCTATGTGATAGACACACAACTTAAGATTTAAAATATTCTAAGCAT	600
                     ************************************************************

2500bp_upstream      ACACTAGTTATATTCAACCTTAAGACTAGATGACCACATATATGCACGGATATAAAGATA	660
GG_promoter          ACACTAGTTATATTCAACCTTAAGACTAGATGACCACATATATGCACGGATATAAAGATA	660
                     ************************************************************

2500bp_upstream      TCCTTTGCCATCTGCTCCAATTGTTGTTATACCACAACTATGTCCTGGAACTCCATCCAC	720
GG_promoter          TCCTTTGCCATCTGCTCCAATTGTTGTTATACCACAACTATGTCCTGGAACTCCATCCAC	720
                     ************************************************************

2500bp_upstream      TTTAGTTCTTGTTCGGTTATTTCAATTTCATGCGGATTGAAGTGTATTGGAAGGGGATGG	780
GG_promoter          TTTAGTTCTTGTTCGGTTATTTCAATTTCATGCGGATTGAAGTGTATTGGAAGGGGATGG	780
                     ************************************************************

2500bp_upstream      ATTTTGACTTGCTATTGATTTAAACCGACTCAATACCACCCAATCCACATGGATTAACGG	840
GG_promoter          ATTTTGACTTGCTATTGATTTAAACCGACTCAATACCACCCAATCCACATGGATTAACGG	840
                     ************************************************************

2500bp_upstream      TGAAATGAACAAGCCCTTAGGGATTGTTCGTTTTAATGTTAATCTATGTGGATCGGGTGT	900
GG_promoter          TGAAATGAACAAGCCCTTAGGGATTGTTCGTTTTAATGTTAATCTATGTGGATCGGGTGT	900
                     ************************************************************

2500bp_upstream      GATTGAATATATCAATAACCTAACAAAGCTTTAGAGAAGCCCGTGGAAAAAAATACATAT	960
GG_promoter          GATTGAATATATCAATAACCTAACAAAGCTTTAGAGAAGCCCGTGGAAAAAAATACATAT	960
                     ************************************************************

2500bp_upstream      ATAACTTATAAAGGAGAGTACACTAGTAGAAAAAGGCTCAAAGCCTGCGGGGACGATATA	1020
GG_promoter          ATAACTTATAAAGGAGAGTACACTAGTAGAAAAAGGCTCAAAGCCTGCGGGGACGATATA	1020
                     ************************************************************

2500bp_upstream      TTTTTACAGACGGATTCGGTTATCCACCGACTGTGTTATTTCTAGTGGCGGTTCCTTAAG	1080
GG_promoter          TTTTTACAGACGGATTCGGTTATCCACCGACTGTGTTATTTCTAGTGgcggttccttaag	1080
                     ************************************************************

2500bp_upstream      AAAACCGCCACTAGAAATCGTATTTCTACTGGCGGTTCCTTAAGAAAACCGCCAGTAGAA	1140
GG_promoter          aaaaccGCCACTAGAA--------------------------------------------	1096
                     ****************

2500bp_upstream      ATCCATGATTTGTAGTGGCGGTTTTCTTAAGGAACCGCCACTAGAAATCATTTTTATCCT	1200
GG_promoter          ---------------------------------------------AATCATTTTTATCCT	1111
                                                                  ***************

2500bp_upstream      TAATTTTTCGAGTTTTTTAAACGGCCTCGTATGACAAAACCACCAAAATAAAAGTTGTAG	1260
GG_promoter          TAATTTTTCGAGTTTTTTAAACGGCCTCGTATGACAAAACCACCAAAATAAAAGTTGTAG	1171
                     ************************************************************

2500bp_upstream      ATCTCTAAAAGTTATGAAACTTTGTAGTTGACAACTTTTTTATTTGAACTCATTTCGGTT	1320
GG_promoter          ATCTCTAAAAGTTATGAAACTTTGTAGTTGACAACTTTTTTATTTGAACTCATTTCGGTT	1231
                     ************************************************************

2500bp_upstream      CTCAAAAATTGAATCTAAGTATGTCAAATTTAAAATTCAAATTTTGCAAACTACCTCGGA	1380
GG_promoter          CTCAAAAATTGAATCTAAGTATGTCAAATTTAAAATTCAAATTTTGCAAACTACCTCGGA	1291
                     ************************************************************

2500bp_upstream      TGAAAAAAGTGTCAAAATAAAAGTTGTAGAACTTCAAAAGTTATTTATCTTTGTAGTTGA	1440
GG_promoter          TGAAAAAAGTGTCAAAATAAAAGTTGTAGAACTTCAAAAGTTATTTATCTTTGTAGTTGA	1351
                     ************************************************************

2500bp_upstream      CAACTTTTTTATTTGAATTCGTTTAGGGTCTCAAATAAGCAATTTACACTCAAATAGTTG	1500
GG_promoter          CAACTTTTTTATTTGAATTCGTTTAGAGTCTCAAATAAGCAATTTACACTCAAATAGTTG	1411
                     ************************** *********************************

2500bp_upstream      TAATATGTGAAGAAAACAACTACAAACTAGACACAAGACATGTCATAGACGGAGTGGTAG	1560
GG_promoter          TAATATGTGAAGAAAACAACTACAAACTAGACACAAGACATGTCATAGACGGAGTGGTAG	1471
                     ************************************************************

2500bp_upstream      GGGAGTGTACGCGCGAGGGTGAGGTCGACGGTTCGAATACTAACAACCGCGTAGCACGCG	1620
GG_promoter          GGGAGTGTACGCGCGAGGGTGAGGTCGACGGTTCGAATACTAACAACCGCGTAGCACGCG	1531
                     ************************************************************

2500bp_upstream      AATTTTGCGCGAAAAATGACGCGACTTGCGATTTTCACTGGCGGTTTTATTAGGTTTTAT	1680
GG_promoter          AATTTTGCGCGAAAAATGACGCGACTTGCGATTTTCACTGGCGGTTTTATTAGGTTTTAT	1591
                     ************************************************************

2500bp_upstream      TACGCCGACCGCCAGTGAAAATCGATTTTCACCGGCGGTCCTCAGTTACCCGCCTGTAAA	1740
GG_promoter          TACGCCGACCGCCAGTGAAAATCGATTTTCACCGGCGGTCCTCAGTTACCCGCCTGTAAA	1651
                     ************************************************************

2500bp_upstream      AATGATGATTTCTACTGGCCCCTAGCACTGGCGGTTACGAAAAACGTCACTATAAATAGG	1800
GG_promoter          AATGATGATTTCTACTGGCCCCTAGCACTGGCGGTTACGAAAAACGTCACTATAAATAGG	1711
                     ************************************************************

2500bp_upstream      TTTACAACCGCCACTATAGAACTTCTCTGTACTAGTGGTATTTTTTCAAATAAAAACAAT	1860
GG_promoter          TTTACAACCGCCACTATAGAACTTCTCTGTACTAGTGGTATTTTTTCAAATAAAAACAAT	1771
                     ************************************************************

2500bp_upstream      TCCTATGCAACGATACAGTCTAACATGCCATTGTTTGTTGTTTCATGGATCTATACAAAC	1920
GG_promoter          TCCTATGCAACGATACAGTCTAACATGCCATTGTTTGTTGTTTCATGGATCTATACAAAC	1831
                     ************************************************************

2500bp_upstream      AAGAAGTTGATTCTAACTCAACATGCATATGTATCACAAAGCTGGAAAGCTAATAACTCT	1980
GG_promoter          AAGAAGTTGATTCTAACTCAACATGCATATGTATCACAAAGCTGGAAAGCTAATAACTCT	1891
                     ************************************************************

2500bp_upstream      AGATGTATGGTAGGGTATATGTACTTAACGTACCTATGTCGTTGCACGTGCAAGCACCCG	2040
GG_promoter          AGATGTATGGTAGGGTATATGTACTTAACGTACCTATGTCGTTGCACGTGCAAGCACCCG	1951
                     ************************************************************

2500bp_upstream      TACCTCACCCCACTACACTACCAATCATTCACAATACATCTCTCTGCTCTTTACTACTAG	2100
GG_promoter          TACCTCACCCCACTACACTACCAATCATTCACAATACATCTCTCTGCTCTTTACTACTAG	2011
                     ************************************************************

2500bp_upstream      CTAGCTAGCTATATAAAGCGGAGCCTGCTAGCTCTCTCTCCCCCATCAGCAGACCACCAC	2160
GG_promoter          CTAGCTAGCTATATAAAGCGGAGCCTGCTAGCTCTCTCTCCCCCATCAGCAGACCACCAC	2071
                     ************************************************************

2500bp_upstream      CCAATCACACCAGCTCTCTCTAGAGCTAGCCCTCTCTTCCTCCAACACTTGTTGATCCCC	2220
GG_promoter          CCAATCACACCAGCTCTCTCTAGAGCTAGCCCTCTCTTCCTCCAACACTTGTTGATCCCC	2131
                     ************************************************************

2500bp_upstream      TCCCATCTCCTCAAGCCTTCTTCACTGAATTTCTGGCCGGTCGATCGTC	2269
GG_promoter          TCCCATCTCCTCAAGCCTTCTTCACTGAATTTCTGGCCGGTCGATCGTC	2180

                     *************************************************


>2500bp_upstream
CGACTAATTTGTTCCGACGGGCTAACTTCCGACGGTTTCGGCTAACTTCCGACGGTTTAAATAACTTCCGAGAGTTTTGCGCTAACTTCCTACGGTTTAGGCCCTCAGAAGTTAAGTATTTTGGTGTAGTGTGTTATATTATAGTTCTCTGAATTTCAAGGCAAGAATTGTATATAACTTCCTAGCGGTACAATCAAAAGAAACGATTCACATTTTTTTCTATAGTCAAAATAATGGCAAATTACATGTTGCCAATGAGGCACTTTATAGGAGAAGTGCCTAAGGCTATGGCCAGCAGATCATACATATGGTCGTGCAAAACACTGTTTACAGAGTATAATTTGAAATGAGAGATATATGAAGATGAGTAACCTGCTAGAGATAGCCTTAAGTGAACTAAATGCTCATCCAAAAAAATTGTGTGAAACCTATATTTATGATTGCTTCTGGAAGACTTTGTTTTTATTATTGCTTTAAAAGAAAGATTTTATTTCATTCCAAGATTAGTACATCGAGTTGATACATGGTCTTAGAATAGTCTCCCAGCAGCCTCTTCTATGTGATAGACACACAACTTAAGATTTAAAATATTCTAAGCATACACTAGTTATATTCAACCTTAAGACTAGATGACCACATATATGCACGGATATAAAGATATCCTTTGCCATCTGCTCCAATTGTTGTTATACCACAACTATGTCCTGGAACTCCATCCACTTTAGTTCTTGTTCGGTTATTTCAATTTCATGCGGATTGAAGTGTATTGGAAGGGGATGGATTTTGACTTGCTATTGATTTAAACCGACTCAATACCACCCAATCCACATGGATTAACGGTGAAATGAACAAGCCCTTAGGGATTGTTCGTTTTAATGTTAATCTATGTGGATCGGGTGTGATTGAATATATCAATAACCTAACAAAGCTTTAGAGAAGCCCGTGGAAAAAAATACATATATAACTTATAAAGGAGAGTACACTAGTAGAAAAAGGCTCAAAGCCTGCGGGGACGATATATTTTTACAGACGGATTCGGTTATCCACCGACTGTGTTATTTCTAGTGGCGGTTCCTTAAGAAAACCGCCACTAGAAATCGTATTTCTACTGGCGGTTCCTTAAGAAAACCGCCAGTAGAAATCCATGATTTGTAGTGGCGGTTTTCTTAAGGAACCGCCACTAGAAATCATTTTTATCCTTAATTTTTCGAGTTTTTTAAACGGCCTCGTATGACAAAACCACCAAAATAAAAGTTGTAGATCTCTAAAAGTTATGAAACTTTGTAGTTGACAACTTTTTTATTTGAACTCATTTCGGTTCTCAAAAATTGAATCTAAGTATGTCAAATTTAAAATTCAAATTTTGCAAACTACCTCGGATGAAAAAAGTGTCAAAATAAAAGTTGTAGAACTTCAAAAGTTATTTATCTTTGTAGTTGACAACTTTTTTATTTGAATTCGTTTAGGGTCTCAAATAAGCAATTTACACTCAAATAGTTGTAATATGTGAAGAAAACAACTACAAACTAGACACAAGACATGTCATAGACGGAGTGGTAGGGGAGTGTACGCGCGAGGGTGAGGTCGACGGTTCGAATACTAACAACCGCGTAGCACGCGAATTTTGCGCGAAAAATGACGCGACTTGCGATTTTCACTGGCGGTTTTATTAGGTTTTATTACGCCGACCGCCAGTGAAAATCGATTTTCACCGGCGGTCCTCAGTTACCCGCCTGTAAAAATGATGATTTCTACTGGCCCCTAGCACTGGCGGTTACGAAAAACGTCACTATAAATAGGTTTACAACCGCCACTATAGAACTTCTCTGTACTAGTGGTATTTTTTCAAATAAAAACAATTCCTATGCAACGATACAGTCTAACATGCCATTGTTTGTTGTTTCATGGATCTATACAAACAAGAAGTTGATTCTAACTCAACATGCATATGTATCACAAAGCTGGAAAGCTAATAACTCTAGATGTATGGTAGGGTATATGTACTTAACGTACCTATGTCGTTGCACGTGCAAGCACCCGTACCTCACCCCACTACACTACCAATCATTCACAATACATCTCTCTGCTCTTTACTACTAGCTAGCTAGCTATATAAAGCGGAGCCTGCTAGCTCTCTCTCCCCCATCAGCAGACCACCACCCAATCACACCAGCTCTCTCTAGAGCTAGCCCTCTCTTCCTCCAACACTTGTTGATCCCCTCCCATCTCCTCAAGCCTTCTTCACTGAATTTCTGGCCGGTCGATCGTC

>GG_promoter
CGACTAATTTGTTCCGACGGGCTAACTTCCGACGGTTTCGGCTAACTTCCGACGGTTTAAATAACTTCCGAGAGTTTTGCGCTAACTTCCTACGGTTTAGGCCCTCAGAAGTTAAGTATTTTGGTGTAGTGTGTTATATTATAGTTCTCTGAATTTCAAGGCAAGAATTGTATATAACTTCCTAGCGGTACAATCAAAAGAAACGATTCACATTTTTTTCTATAGTCAAAATAATGGCAAATTACATGTTGCCAATGAGGCACTTTATAGGAGAAGTGCCTAAGGCTATGGCCAGCAGATCATACATATGGTCGTGCAAAACACTGTTTACAGAGTATAATTTGAAATGAGAGATATATGAAGATGAGTAACCTGCTAGAGATAGCCTTAAGTGAACTAAATGCTCATCCAAAAAAATTGTGTGAAACCTATATTTATGATTGCTTCTGGATGACTTTGTTTTTATTATTGCTTTAAAAGAAAGATTTTATTTCATTCCAAGATTAGTACATCGAGTTGATACATGGTCTTAGAATAGTCTCCCAGCAGCCTCTTCTATGTGATAGACACACAACTTAAGATTTAAAATATTCTAAGCATACACTAGTTATATTCAACCTTAAGACTAGATGACCACATATATGCACGGATATAAAGATATCCTTTGCCATCTGCTCCAATTGTTGTTATACCACAACTATGTCCTGGAACTCCATCCACTTTAGTTCTTGTTCGGTTATTTCAATTTCATGCGGATTGAAGTGTATTGGAAGGGGATGGATTTTGACTTGCTATTGATTTAAACCGACTCAATACCACCCAATCCACATGGATTAACGGTGAAATGAACAAGCCCTTAGGGATTGTTCGTTTTAATGTTAATCTATGTGGATCGGGTGTGATTGAATATATCAATAACCTAACAAAGCTTTAGAGAAGCCCGTGGAAAAAAATACATATATAACTTATAAAGGAGAGTACACTAGTAGAAAAAGGCTCAAAGCCTGCGGGGACGATATATTTTTACAGACGGATTCGGTTATCCACCGACTGTGTTATTTCTAGTGgcggttccttaagaaaaccGCCACTAGAAAATCATTTTTATCCTTAATTTTTCGAGTTTTTTAAACGGCCTCGTATGACAAAACCACCAAAATAAAAGTTGTAGATCTCTAAAAGTTATGAAACTTTGTAGTTGACAACTTTTTTATTTGAACTCATTTCGGTTCTCAAAAATTGAATCTAAGTATGTCAAATTTAAAATTCAAATTTTGCAAACTACCTCGGATGAAAAAAGTGTCAAAATAAAAGTTGTAGAACTTCAAAAGTTATTTATCTTTGTAGTTGACAACTTTTTTATTTGAATTCGTTTAGAGTCTCAAATAAGCAATTTACACTCAAATAGTTGTAATATGTGAAGAAAACAACTACAAACTAGACACAAGACATGTCATAGACGGAGTGGTAGGGGAGTGTACGCGCGAGGGTGAGGTCGACGGTTCGAATACTAACAACCGCGTAGCACGCGAATTTTGCGCGAAAAATGACGCGACTTGCGATTTTCACTGGCGGTTTTATTAGGTTTTATTACGCCGACCGCCAGTGAAAATCGATTTTCACCGGCGGTCCTCAGTTACCCGCCTGTAAAAATGATGATTTCTACTGGCCCCTAGCACTGGCGGTTACGAAAAACGTCACTATAAATAGGTTTACAACCGCCACTATAGAACTTCTCTGTACTAGTGGTATTTTTTCAAATAAAAACAATTCCTATGCAACGATACAGTCTAACATGCCATTGTTTGTTGTTTCATGGATCTATACAAACAAGAAGTTGATTCTAACTCAACATGCATATGTATCACAAAGCTGGAAAGCTAATAACTCTAGATGTATGGTAGGGTATATGTACTTAACGTACCTATGTCGTTGCACGTGCAAGCACCCGTACCTCACCCCACTACACTACCAATCATTCACAATACATCTCTCTGCTCTTTACTACTAGCTAGCTAGCTATATAAAGCGGAGCCTGCTAGCTCTCTCTCCCCCATCAGCAGACCACCACCCAATCACACCAGCTCTCTCTAGAGCTAGCCCTCTCTTCCTCCAACACTTGTTGATCCCCTCCCATCTCCTCAAGCCTTCTTCACTGAATTTCTGGCCGGTCGATCGTC
```
